# Cholinergic-dependent dopamine signals in mouse dorsomedial striatum are regulated by frontal but not sensory cortices

**DOI:** 10.1101/2025.09.30.679538

**Authors:** Hannah C. Goldbach, Rachele Rimondini, Evan S. Swanson, Jung Hoon Shin, Michael E. Authement, Lucy G. Anderson, Han Bin Kwon, Ron Paletzki, Charles R. Gerfen, Linda M. Amarante, Richard J. Krauzlis, Veronica A. Alvarez

## Abstract

Everyday decisions depend on linking sensory stimuli with actions and outcomes. The striatum supports these sensorimotor associations through dopamine-dependent plasticity. Thus, the timing and magnitude of dopamine release is critical for learning. Recent work has characterized a local striatal microcircuit in which cholinergic interneurons (CINs) modulate dopamine release via acetylcholine activation of nicotinic receptors on dopamine axons. Here, we show that visual stimuli evoke dopamine responses in the dorsomedial striatum through this cholinergic-dependent mechanism. Using anatomical and functional methods to identify which pathways elicit these signals, we found that visual and auditory cortices that project to the dorsomedial striatum lack robust connectivity to CINs and were unable to drive cholinergic-dependent dopamine release. In contrast, frontal cortical regions, including the prelimbic and anterior cingulate cortices, strongly recruited CINs and acetylcholine, producing robust dopamine release both *ex vivo* and *in vivo*. These frontal corticostriatal projections are activated by visual stimuli, representing a possible pathway by which visual information reaches the dorsomedial striatum to evoke dopamine. These findings reveal a fundamental distinction between sensory and frontal cortical inputs to the striatum, demonstrating that only the latter evoke cholinergic-dependent dopamine signaling. This work establishes a framework for understanding how cortical circuits shape striatal dopamine to support reinforcement learning.

## Introduction

Sensory input provides a nearly constant flux of information that guides our decisions about actions, based on expected outcomes. However, the complex pathways through which sensory information drives action remain unclear. The dorsomedial striatum is one brain area known to play a role in this process^1–4^. The dorsomedial striatum receives glutamatergic input from sensory cortex, frontal cortex, and thalamic nuclei, as well as dopaminergic input from the midbrain. It is this integration of contextual sensory information with expected outcomes, signaled largely by dopamine, that enables dorsomedial striatum’s function in guiding goal-directed behaviors, sensory learning, and attention. Historically, most mechanistic understanding has focused on how striatal medium spiny neurons (MSNs), which comprise the main output pathways of the striatum, are driven by glutamatergic inputs and modulated by dopamine^5,6^. The temporal alignment of these glutamatergic and dopamine signals is crucial as it determines the rules of synaptic plasticity, promoting either long-term potentiation or depression of selective inputs^7–9^ to reshape functional connectivity and promote the associative learning of sensory cues and motor actions^10–13^.

Midbrain-originating action potentials have long been thought to have sole control over the timing of striatal dopamine signals, so much so that midbrain neuron activity is commonly interpreted as a proxy for dopamine release in the striatum. However, an alternative mechanism has recently been identified, in which acetylcholine released by cholinergic interneurons (CINs) can act locally within the striatum to regulate dopaminergic axonal excitability and dopamine release through nicotinic acetylcholine receptors (nAChRs)^14–21^, independently of soma-generated action potentials^22,23^. This local regulatory mechanism does not reflect the activity of CINs alone. Instead, CIN-evoked dopamine appears to act as a coincidence detector by requiring synchronized recruitment by the glutamatergic inputs arriving from the cortex and thalamus^17^.

Striatal CINs produce a characteristic response to novel or salient stimuli, which can strengthen as an animal learns to associate those stimuli with actions and rewards. This has been observed both in non-human primates^24–27^ and, more recently, in mice^28,29^. The recently identified ability of CINs to locally regulate dopamine opens new possibilities about the neural circuits underlying these behavioral findings. Theoretically, cortical and thalamic projections to the striatum that recruit CINs could regulate the temporal and spatial distribution of dopamine signals to shape behavior and learned associations with sensory stimuli. In support of a physiological role for this local mechanism, a recent study demonstrated that cholinergic control over dopamine in the ventral striatum contributes to effortful behavior^30^. Recent experiments have shown that motor and somatosensory cortical inputs, as well as thalamic inputs, synapse onto MSNs with more strength than they do onto CINs^31,32^. Yet we do not know which glutamatergic inputs drive striatal CINs to provide cholinergic control over dopamine, nor how it compares across subregions of the striatum. Recent *in vivo* evidence suggests that CIN responses to sensory cues are shaped by convergence of top-down inputs from the cortex with bottom-up inputs from the thalamus and midbrain^33^. Therefore, CINs might play an integrative role in the formation of associations for sensory-guided actions, whereby brain regions with functional connections to CINs can synchronize acetylcholine release to gain temporal and spatial control over dopamine transmission.

These findings raise new fundamental questions about which inputs to the dorsomedial striatum recruit CINs to regulate dopamine transmission. Motor cortex^18^, prelimbic cortex^20^, and some thalamic nuclei^18,34^ can evoke striatal dopamine through recruitment of CINs. However, it is unknown whether sensory cortical areas, which project robustly to the dorsomedial striatum can also recruit CINs directly. Specifically, if sensory cortices could also recruit CINs and control the timing of dopamine signals, this would allow direct control over action selection and the learned associations between stimuli and rewards. In contrast, if the ability to drive striatal dopamine through CINs is unique to a few areas—such as prefrontal cortex and thalamus—it would indicate a larger role for feedforward pathways through the frontal cortex and sensory thalamic nuclei in sensory-guided learning.

Here we show that, in fact, salient visual stimuli elicit striatal dopamine in part via a cholinergic-dependent mechanism. We identify specific cortical areas and striatal cell types involved in driving cholinergic-dependent dopamine release in the dorsomedial striatum. *In vivo* fiber photometry and *ex vivo* recordings revealed that prefrontal areas connect far more heavily with CINs in dorsomedial striatum than cortical sensory areas such as visual and auditory cortices, despite their prominent projections to striatal medium spiny neurons in dorsomedial striatum. Cortical sensory areas failed to drive cholinergic-dependent dopamine in dorsomedial striatum and formed only rare and weak synaptic contacts with CINs, as validated by anatomical and functional analysis. Lastly, we confirmed that visual stimuli drive *in vivo* time-locked calcium responses in striatal-projecting frontal cortical axons and stimulation of these corticostriatal projections recruits CINs to drive acetylcholine and dopamine release in the dorsomedial striatum.

Through this combination of experimental approaches, our findings establish that frontal cortices, but not sensory cortices, can play a prominent role in controlling striatal CINs and modulating striatal dopamine. These results complement other findings demonstrating the importance of frontal cortices in sensorimotor learning and dopamine transmission, and highlight the importance of these new circuit details as an unexplored avenue for understanding the etiology of dopamine-related disorders.

## Results

### Salient visual stimuli elicit striatal dopamine through both midbrain and local cholinergic mechanisms

We used fiber photometry to survey dopamine release and dopamine neuron activity in response to salient visual stimuli (**Fig. 1a,e**, Methods). DAT-cre mice expressing cre-dependent GCaMP8s in midbrain dopaminergic neurons were implanted with two fiber optic cannulae: one in the midbrain to measure somatic dopamine neuron activity, and another in dorsomedial striatum (DMS) to measure axonal dopamine neuron activity (**Fig. 1b-c**). A separate group of mice, expressing the dopamine sensor dLight1.3b in DMS were implanted with fiber optic cannulae in the DMS to measure striatal dopamine (**Fig 1d**). All mice were implanted with headframe and habituation to the headfixed chamber. Mice passively experienced visual stimuli (LED light, 5.9 cd/m^2^, 500 ms duration) delivered at random intervals averaging 60 s. Calcium responses time-locked to visual stimuli were recorded from both the somas and the axons of the dopamine neurons (**Fig. 1f,h**; mean z-score in somas 1.58 [1.28 - 1.88 bootstrap 95% CI], n = 5 animals; mean in axons 1.90 [1.53 - 2.39], n = 7 animals). Within the DMS, these same visual stimuli elicited a robust dopamine signal time-locked to stimulus onset, measured as large transients of dLight1.3b fluorescence (**Fig. 1g-h**; mean z-score 3.70 [1.53 - 2.39], n = 25 animals; significant effect of sensor and recording location p = 0.04, one-way ANOVA).

**Figure 1:**
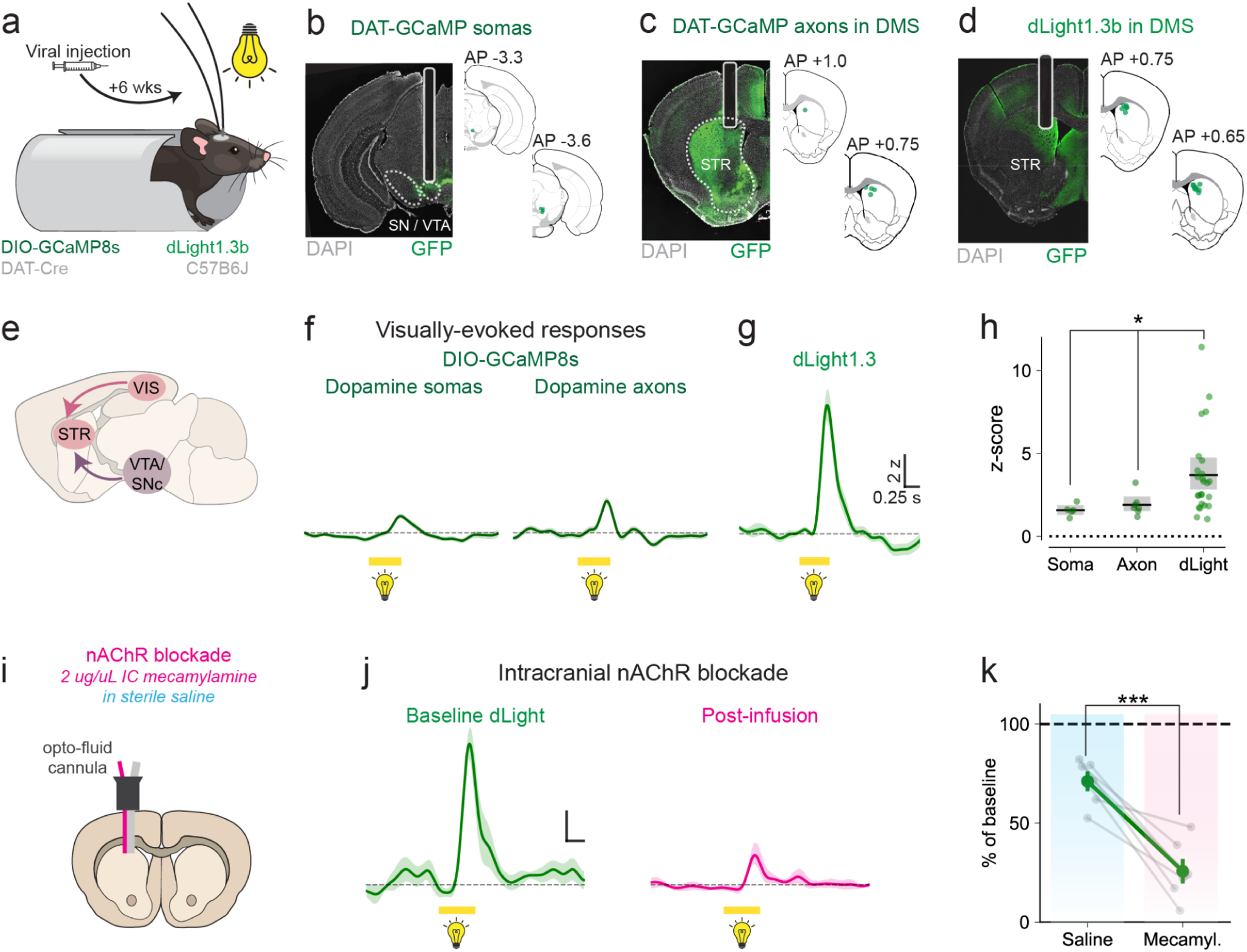
Salient sensory stimuli trigger nAChR-dependent striatal dopamine. **a.** Experimental setup of head-fixed mice presented with white light (500 ms, 5.9 cd/m^2^). Mice were held in a dark sound-attenuating box and stimuli presentation spaced roughly every 60 seconds. Intracranial surgeries were carried out at least 6 weeks in advance. **b-d.** Representative images of immuno-fluorescence from DAT-IRES-Cre mice expressing GCaMP8s in midbrain dopamine neurons with fiber implants in midbrain and DMS (b,c); or wildtype mice expressing dLight1.3b in DMS with fibers implants in DMS (d). **e.** Diagram depicts a canonical pathway (purple) for dopamine signals in the striatum generated from ventral tegmental area (VTA) and substantia nigra compacta (SNc). Additional putative pathway (pink) for generation of nAChR-dependent striatal dopamine in response to visual stimuli. **f-g.** Average example traces of visual stimulus evoked calcium signals (f) from midbrain fibers (dopamine neuron somas) and DMS fiber (axons) and dLight1.3 dopamine signals in DMS. **h** Maximum z-scored for evoked GCaMP8 (soma n = 5, axon n = 7) and dLight1.3 (n = 25) responses to light cue presentation. Each dot represents a mouse, black lines represent medians, and gray bars represent the bootstrap 95% CI. **i.** A subset of dLight-expressing mice were implanted with optofluid cannula and received intracranial infusions of saline (vehicle) or mecamylamine (0.2 ug / uL). **j.** Single session traces from a representative animal showing the visually-evoked dopamine response before (baseline) and after (magenta) intracranial infusion of mecamylamine. **k.** Across-animal effects of intracranial saline and mecamylamine on visually-evoked dopamine responses. Amplitudes were normalized to baseline responses before infusion (n = 7 mice). Gray dots are individual animals and green dots represent the mean +/– sem. For all traces, line and shade represent mean z-score +/- sem.

Given recent reports that striatal acetylcholine may amplify or regulate striatal dopamine^35–38,21^, we explored whether these visually-evoked dopamine signals were dependent on nicotinic receptor activation. We first tested the effect of systemic administration of mecamylamine, a non-competitive nicotinic receptor antagonist. We found that mecamylamine (10 mg/kg, i.p.) reduced visual-evoked dopamine signals to 41 ± 4.8% of baseline, while sparing the somatic calcium responses in dopamine neurons (**Suppl. Fig. 1**). These results from systemic drug administration hinted at a cholinergic contribution but could not distinguish between a striatal site of action and peripheral effects. To directly test the effects of mecamylamine in the striatum, we repeated the dopamine sensor experiments using an opto-fluid cannula that allows for intracranial infusion within the recorded area of mecamylamine (2 ug / uL) or vehicle (saline) (**Fig. 1i**). The effects were even larger – intracranial mecamylamine significantly decreased visually-evoked dopamine responses to 26 ± 5% **(Fig. 1j-k**; n = 7 animals; treatment p < 0.0001, one-way repeated-measures ANOVA).

We reasoned that a visually evoked, cholinergic-dependent dopamine signal would require the recruitment of striatal cholinergic interneurons (CINs) by visual stimuli. Striatal CINs have been shown to respond to salient sensory stimuli in a series of classical electrophysiology studies done in non-human primates^24,25^. To assess whether the visual stimulus drove striatal CINs and evoked local ACh release in the mouse DMS, we expressed unilateral GCaMP8 in CINs and/or unilateral GRAB-ACh3.0, an acetylcholine sensor (**Fig. 2a-c, Table S1**). Calcium signals from CINs increased in response to visual stimuli (**Fig. 2d-e**; mean 1.84 z [1.58 - 2.08], n = 12 animals), confirming a visual drive. Furthermore, visual stimuli increased acetylcholine in DMS, as measured with GRAB-ACh3.0 (**Fig. 2g-h**; mean 3.30 z [2.31 - 4.31], n =7 animals). Note that both cue-evoked responses were notably multiphasic and heterogeneous, consisting of one, two, and sometimes even three phases (**Suppl. Fig. 2**), in line with recent reports^39^. Quantification was limited to the first phase of the response.

**Figure 2:**
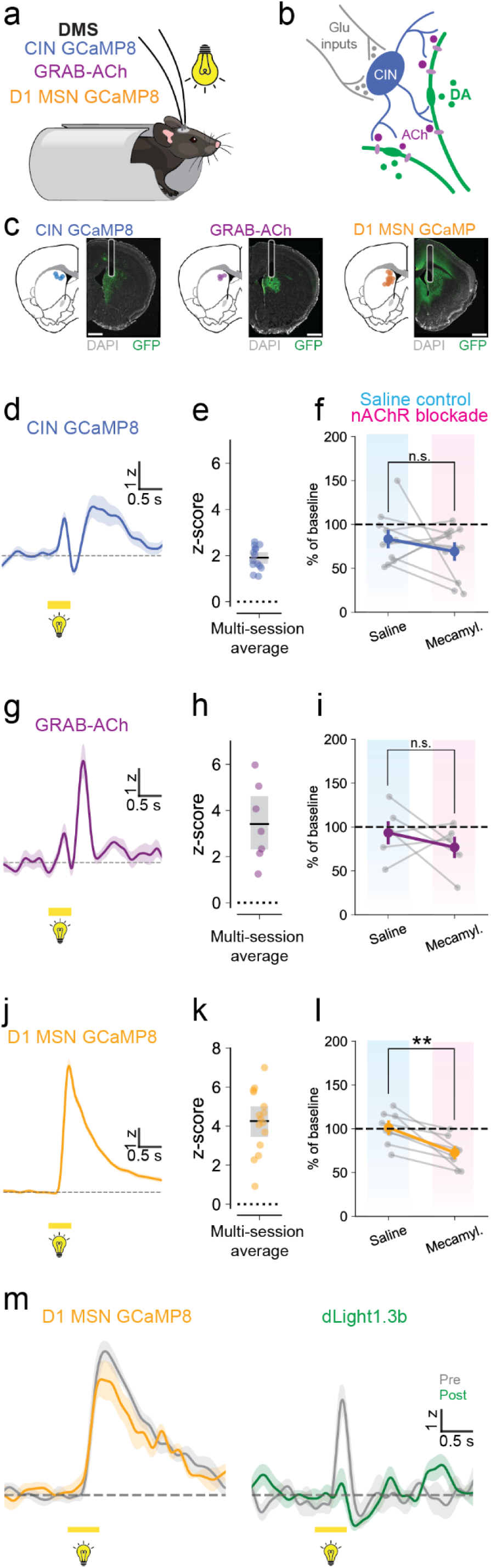
Visual stimuli recruit striatal CINs and drive acetylcholine responses. **a** Headfixed setup for light stimulus presentation (5.9 cd/m^2^, 500 ms). All mice underwent surgery at least 6 weeks in advance and received fiber implants in DMS. ChAT-IRES-Cre and Drd1-Cre mice were used to express GCaMP in CINs or D1-MSNs, respectively. Wildtype C57Bl-6J or ChAT-IRES-cre mice expressed GRAB-ACh in the dorsomedial striatum. **b.** Diagram of the striatal microcircuitry possibly involved: cholinergic interneurons (blue) are recruited by glutamatergic inputs to the striatum (gray) and release acetylcholine (purple), which activates nAChRs on DA axons (green) to produce local dopamine release. **c.** Fiber placements and histology from representative mice expressing GCaMP8 in CINs, GRAB-ACh, or GCaMP8 in D1-MSNs. Scale bars are 1 mm. **d, g, j.** Single-animal traces of GCaMP responses in CINs (d), GRAB-ACh (g), and GCaMP responses in D1-MSNs (j) aligned to light onset. Lines and shades are mean z-score +/- sem. **e, h, k.** Maximum z-score of visual-evoked responses for GCaMP in CINs (e, n= 12 mice), GRAB-ACh (h, n = 7 mice), and GCaMP in D1-MSNs (k, n = 9 mice). Each dot represents one animal.. Black lines represent means, and gray bars represent bootstrapped 95% CI. **f, i, l.** Normalized effect of saline and nAChR antagonist mecamylamine (10 mg/kg, i.p.) on in vivo responses to light onset in mice expressing GCaMP in CINs (f, n = 11 mice), GRAB-ACh (i, n = 6 mice) or GCaMP in D1-MSNs (k, n = 8 mice). Gray symbols and lines are paired data from individual hemispheres pre- and post-administration of saline (blue) or mecamylamine (pink). Black lines and shaded bars represent the median and bootstrapped 95% CI. **m.** Representative traces show visual-evoked responses from an animal expressing GCaMP8 in D1-MSNs in one hemisphere (left) and dLight1.3b in the other hemisphere (right). GCaMP signals in D1-MSN were only slightly reduced by mecamylamine administration, while dLight1.3b signals were dramatically reduced. All traces represent mean z-score +/- sem.

In contrast to its effects on dopamine signals, systemic mecamylamine (10 mg/kg, i.p) had no effect on visually-evoked calcium signals in CINs (**Fig. 2f**; n = 11 animals; p = 0.21, one-way repeated-measures ANOVA) nor acetylcholine release (**Fig. 2i**; n = 6 animals; p = 0.43, one-way repeated-measures ANOVA). This insensitivity to nicotinic receptor blockers is consistent with the candidate circuit mechanism (**Fig. 2b**), since CIN activity and acetylcholine release lie upstream of the proposed site of action of nACh receptors on the dopamine axon terminals. It also further rules out peripheral actions or even central action on other brain circuits of systemic mecamylamine administration.

As a control, we also confirmed that D1 receptor-expressing MSNs robustly respond to the visual stimulus using the same sensor GCaMP8s (**Fig. 2j-k**; mean 4.06 z [2.96 - 5.04] n = 9 animals). Calcium signals in D1 MSNs persisted after systemic mecamylamine, though with a small decrease (**Fig. 2l-m**; n = 8 animals; p = 0.0035, one-way repeated-measures ANOVA), possibly due to the blunting of the dopamine response by mecamylamine. Together, the fact that cue-evoked responses in D1, CIN, and acetylcholine were largely insensitive to nAChR blocker is evidence that visual information still reached the striatum and argues against a generalized impairment in visual pathways. Despite the presence of a preserved visual drive, we confirmed a dramatic decrease in visually-evoked dopamine responses during systemic mecamylamine administration (**Fig. 2m, right**), providing further evidence that activation of nicotinic ACh receptors is a key element underlying the sensory stimuli-evoked dopamine responses.

We also tested auditory responses to determine whether the findings generalized to other sensory modalities. In this anterior portion of the dorsomedial striatum, brief tone presentations (7.5 KHz at 70 dB for 500 ms) failed to produce any reliable responses in dopamine somas or axons, despite robust responses in somas to waterdrop reward delivery (**Suppl. Fig. 3**). Similarly, tones failed to produce consistent dopamine response (**Suppl. Fig. 4**). Auditory cues also failed to trigger acetylcholine transients nor recruit CINs in this anterior DMS (**Suppl. Fig. 4**). Thus, the sensory-evoked striatal dopamine responses in the anterior DMS may be relatively selective for visual stimuli.

### Unlike frontal cortical areas, primary sensory areas do not synapse onto dorsomedialstriatal CINs

Next, we asked which striatal inputs could drive striatal CINs to promote local acetylcholine and, potentially, dopamine release. Primary sensory areas (visual, somatosensory, and auditory cortex) are known to project to the dorsal striatum. Though known to synapse with MSNs, it is not yet known whether they also synapse onto CINs. To address this, we used cell-type-specific monosynaptic retrograde rabies tracing to widely survey the frequency of cortical connections to striatal CINs in the dorsal striatum. We virally expressed cre-dependent TVA receptors and rabies glycoprotein in striatal CINs of ChAT-cre mice, which allowed a modified glycoprotein-deleted rabies virus to selectively infect cre-expressing CINs (“starter cells”) and label neurons that synapse onto them (**Fig. 3a-b; Table S1; Suppl. Fig. 5; Methods)**. The majority of labeled projection cells were identified in the cortex (49-54%), although labeled cells were also found in the thalamus (5-9%), midbrain (1-2%), and locally within basal ganglia (38-44%; **Fig. 3f, insets**)

**Figure 3.**
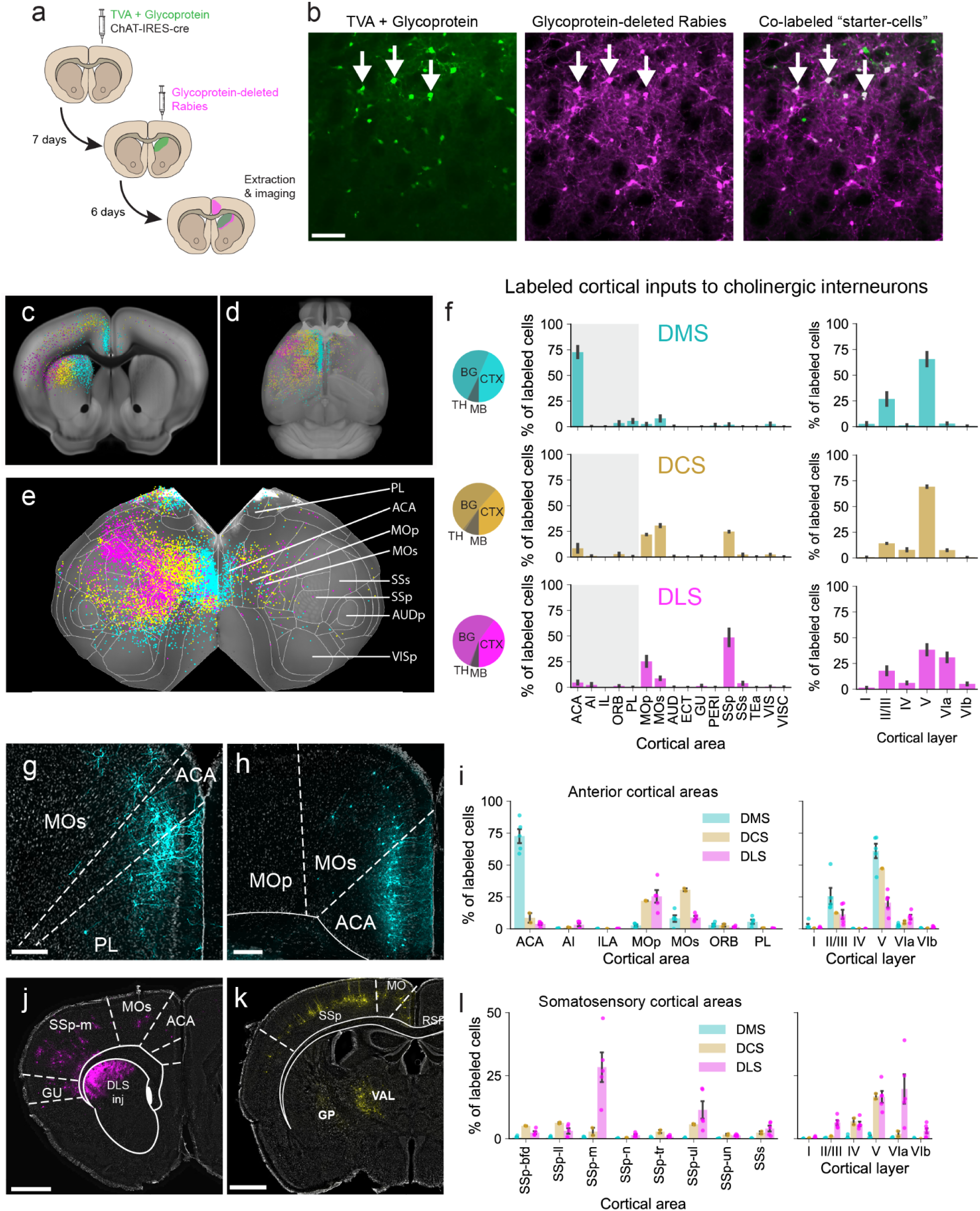
Retrograde rabies tracing demonstrates a widespread anterior bias towards frontal cortical connectivity with striatal cholinergic interneurons. **a.** Experimental timeline for injecting ChAT^IRES-cre^ mice with cre-dependent TVA rabies receptor + glycoprotein, followed by a modified glycoprotein-deleted retrograde rabies vector to label CIN projecting cells. **b.** Images of a DMS injection area showing cells expressing TVA-glycoprotein (left, green) and glycoprotein-deleted rabies (middle, magenta). Overlaid images (right) show co-labeled cells that serve as “starter” cells (white). **c, d, e.** Reconstructions showing CIN-projecting cortical cells in coronal view (c), whole-brain view (d), and flattened brain view (e) from injections in DMS (cyan), DCS (yellow), and DLS (magenta). **f.** Quantifications of CIN-projecting labeled cortical cells per area (left) and per cortical layer (right). Plots show averages of all labeled cortical cells for injection sites in DMS (n = 5 mice), DCS (n = 2 mice), and DLS (n = 5 mice). ACA, anterior cingulate area; PL, prelimbic; MOp, primary motor; MOs, secondary motor; SSp, primary somatosensory. See Table S2 for other anatomical nomenclature. Inset pie charts represent percentages of labeled cells in cortex (CTX), basal ganglia (BG), thalamus (TH), and midbrain (MB). **g-h, j-k.** Representative images of retrogradely labeled cortical neurons projecting to striatal CINs in DMS (g-h), to CINs in DCS (j), and to CINs in DLS (k) in coronal sections. Inset scale bars are 250 μm. **i, l.** Quantifications of retrogradely labeled cortical cells per subregion (left) and layer (right) expressed as a percentage of total labeled cortical cells for frontal cortical areas (i) and somatosensory areas (l).

Labeling across the medial to lateral axis in the cortex mirrored injection sites across the same axis in the striatum. Injections in medial, central, and lateral dorsal striatum produced labeling in medial frontal areas, motor cortices, and somatosensory cortex, respectively (**Fig. 3c-f**). Most of this labeling was found in anterior cortical areas, with only sparsely labeled cells in more posterior areas, including primary visual cortex (VISp) and primary auditory cortex (AUDp, **Fig. 3c-f**).

Injections in the DMS led to robust labeling in the anterior cingulate area (ACA) with only sparsely labeled cells in sensory cortices (**Fig. 3f**, left). The majority of cells labeled in the cortex were located in layer 5 (66% of labeled cortical cells), although a smaller percentage were located in layers 2/3 (27%) (**Fig. 3f**, right), as expected for striatal projecting neurons^40–45^. Of all cortical cells labeled per mouse, an average of 73% were located in ACA (**Fig. 3g-i**), with additional labeling in the secondary motor cortex (8% of cortical cells, MOs) and prelimbic cortex (6%, PL) (**Fig. 3g, i**). Very few labeled cells were located in primary somatosensory cortex (SSp, 2%), VISp (3%), or AUDp (0.4%) (**Fig. 3f, Suppl. Fig. 2**). Injections in the dorsal central striatum (DCS) led to fairly even labeling across primary motor cortex (MOp, 22% of cortical cells), secondary motor cortex (MOs, 31%), and SSp (25%). Meanwhile, injections in the dorsal lateral striatum (DLS) led to labeling in the heaviest expression in SSp (49% of labeled cortical cells) and lesser expression in MOp (25%) and MOs (9%, **Fig. 3f,k**). Again, the majority of these cells in both DCS and DLS were located in layer 5 (69% and 38%, respectively), with smaller percentages in layer 6 (8%, and 31%) and layers 2/3 (14% and 18%) (**Fig. 3f**).

SSp contains seven topographic sub-regions, each representing a part of the body. When we divided the DLS cell counts further into these sub-regions, we found that roughly half of the labeled SSp cells were located in the mouth/jaw area (SSp-m, 28% out of a total 49%), and one quarter of SSp cells were in the upper limb (SSp-ul, 11% out of 49%, **Fig. 3k-l**). The barrel field of somatosensory cortex (SSp-bfd) contained only 2% of the labeled cells across the cortex (**Fig. 3k-l**). In comparison, 3% of all labeled cells were also found across the visual cortex, which also consists of several sub-regions. VISp contained only 0.6% of labeled cells, while secondary visual areas (VISa, VISal, VISam, VISl, VISli, VISpm, and VISrl) contained less than 1% (0.05 - 0.7%) of labeled cells (**Suppl. Fig. 2**). The auditory cortex contained even fewer labeled cells than the visual cortex, with only 0.03 to 0.2% of cells in each auditory sub-region (**Suppl. Fig. 2**).

These results demonstrate that only a handful of medial frontal cortical areas synapse frequently with striatal CINs in the dorsomedial striatum.

### Frontal areas form reliable functional connections with striatal CINs, while primary sensory areas do not

The retrograde rabies tracing identified specific cortical areas with a significant number of cells forming synapses onto striatal CINs, but it did not inform us about the strength of those synapses. We therefore sought to functionally validate the rabies results using *ex vivo* whole-cell patch-clamp electrophysiology to record excitatory postsynaptic currents (EPSCs) from striatal MSNs and CINs as we optogenetically stimulated different cortical inputs. Mice were injected with ChR2-eYFP or ChrimsonR-tdTomato in either the anterior cingulate cortex, prelimbic cortex, primary somatosensory, visual, or auditory cortex (Methods; **Fig. 4a-c**). All CIN recordings were conducted using ChAT-tdTomato mice to aid in cell identification, while MSN recordings were conducted using ChAT-tdTomato mice or other cre-recombinase lines on C57B6/J backgrounds (**Methods**).

**Figure 4:**
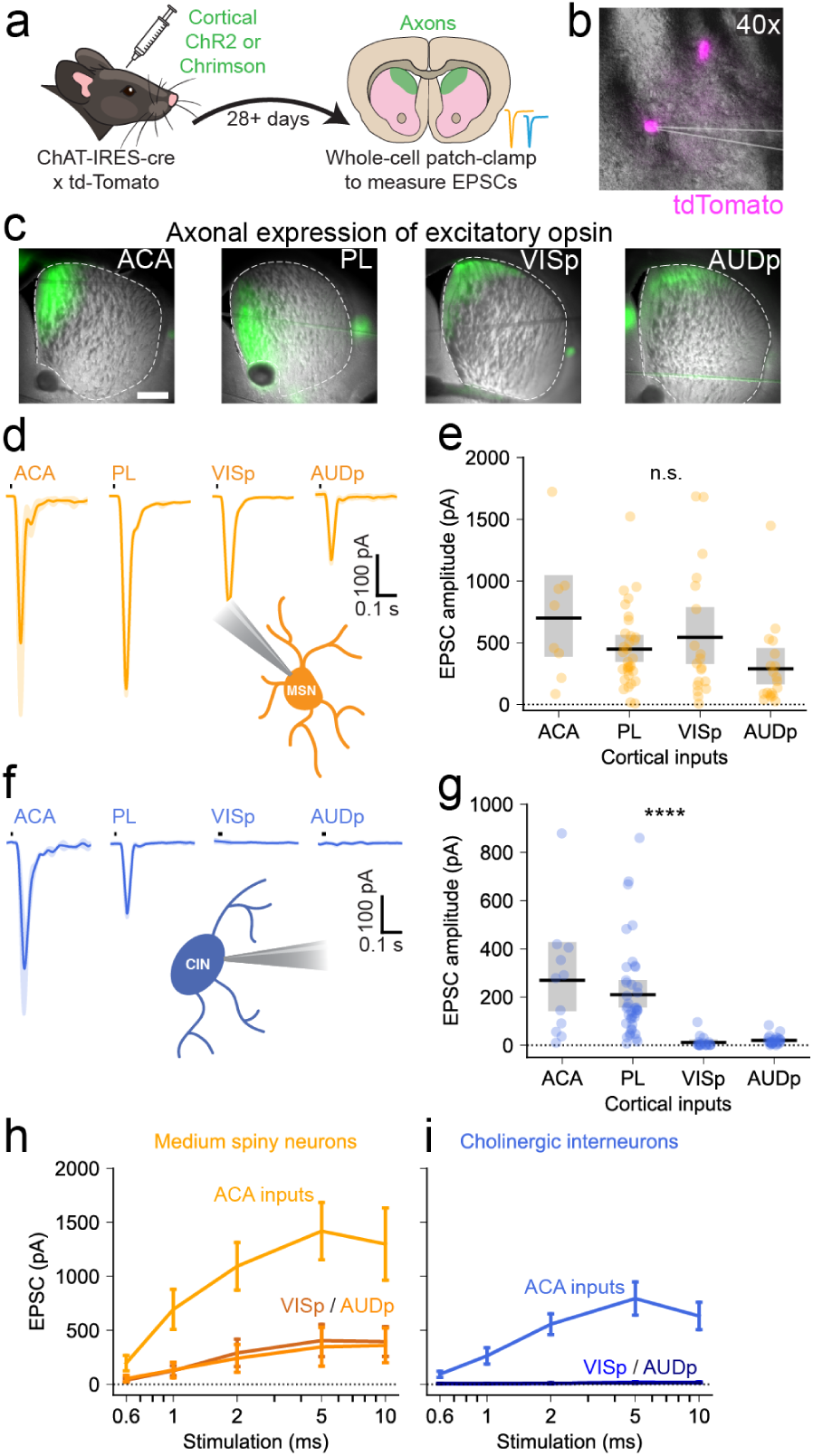
Frontal, but not visual or auditory, cortical areas form reliable functional connections with striatal CIN. **a.** Mice with labeled CINs (ChAT-IRES-Cre x tdTomato) were injected with either ChR2-eYFP or ChrimsonR-tdTomato in anterior cingulate (ACA), prelimbic (PL), visual cortex (VIS) or auditory cortex (AUD). Brains were extracted and sliced for ex vivo whole-cell patch clamp electrophysiology following a minimum four-week incubation. **b.** Visualization of genetically-encoded tdTomato expression in striatal CINs, as viewed on the electrophysiology rig. **c.** Representative images show green fluorescent axonal projections expressing excitatory opsins in DMS, following injections in anterior cingulate (ACA), prelimbic (PL), visual (VISp), or auditory (AUDp) cortices. Scale bar is 500 μm. **d.** Representative traces of excitatory postsynaptic currents (EPSC) recorded from medium spiny neurons (MSNs) in DMS in response to optogenetic stimulation of cortical projections. **e.** Average MSN EPSC amplitudes evoked via optogenetic stimulation of different cortical area projections (ACA n = 11/3; PL n = 41/11; VIS n = 18/7; AUD n= 25/7 - slices/mice). **f.** Representative EPSC traces recorded from CINs in DMS in response to optogenetic stimulation of cortical area projections. **g.** Average CIN EPSC amplitudes evoked via optogenetic stimulation of cortical area projections. (ACA n = 8/3; PL n = 32/8; VIS n = 19/5; AUD n= 19/5 - slices/mice) **h.** Input-output curves showing EPSC amplitudes recorded from MSNs (orange) and CINs (blue) in response to increasing duration of optogenetic stimulation of corticostriatal projections. All traces are mean and shaded sem. For group plots, symbols represent data from individual cells; black lines and shaded bars represent median and bootstrap 95% CI.

Coronal brain slices showed distinctive patterns of labeled corticostriatal projections in the dorsomedial or dorsolateral striatum, depending on the targeted cortical areas (**Fig. 4c**). CIN and MSN recordings were made from the striatal region receiving labeled projections. 405 nm blue light (1.3 mW) and 590 nm orange light (0.7 mW) were used to optogenetically excite these terminals, as a subset of these animals expressed both opsins (see below, **Suppl. Fig. 6**). When recording from MSNs, optogenetic activation (1 ms) of ACA or PL inputs produced large excitatory postsynaptic currents (**Fig. 4d-e**; ACA: mean 699.4 [382.9 - 1062.8 bootstrap 95% CI] pA, n = 8 cells from 3 animals; PL: 448.1 [344.9 - 560.7] pA, n = 32 cells from 9 animals). In line with previous findings, sensory areas also produced measurable EPSCs in MSNs (**Fig. 4d-e**; VISp: 544.0 [327.2 - 792.4] pA, n = 19 cells from 5 animals; AUDp: 289.1 [165.4 - 452.6] pA, n = 19 cells from 5 animals). The tested cortical regions all formed robust synaptic connections with MSNs and we found no significant difference in the mean responses (p = 0.12, one-way ANOVA).

When recording from CINs, however, we only saw large EPSCs in response to a 10 ms stimulation of ACA or PL (**Fig. 4f-g**; ACA: 269.6 [143.6 - 423.3] pA, n = 11 from 3 animals; PL: 209.4 [155.2 - 270.6] pA, n = 41 cells from 12 animals). Stimulation of inputs from primary sensory areas (10 ms) consistently produced little to no response in CINs (**Fig. 4f-g**; VISp: 12.2 [3.8 - 24.2] pA, n = 18 cells from 7 animals; AUDp: 20.2 [13.8 - 28.0] pA, n = 25 cells from 7 animals). Comparing the strength of the synaptic connections to CINs across the cortical regions revealed significant differences between ACA/Pl and the other sensory areas (p < 0.0001, one-way ANOVA). We explored the responses in CIN and MSNs by varying the duration of the stimulation (**Fig. 4h-i**). When considering the sensory areas, a 2-5 ms stimulation was sufficient to saturate the responses recorded from MSNs (**Fig. 4h**). Meanwhile, a 10 ms stimulation of sensory terminals failed to produce any response in CINs (**Fig. 4i**).

As an additional test of the selective connectivity from sensory cortices to MSNs and not CINs, a cohort of mice simultaneously expressed ChrimsonR in a primary sensory area and ChR2 in the frontal cortex (ACA / PL), which served as an internal control. This allowed us to compare how the same cell (a single MSN or single CIN) responded to two different cortical inputs (**Suppl. Fig. 6**). The paired data confirmed the findings that single CINs responded strongly to inputs from PL, but not any primary sensory area. This data also confirmed that single MSNs receive inputs from both frontal and sensory cortices. This experiment provided the benefit of paired data and internal control, but also restricted our recordings spatially to places with overlapping expression.

Our rabies tracing results showed that the primary somatosensory cortex (SSp) connects with CINs in DLS. We therefore repeated these electrophysiology experiments in DLS with stimulation of SSp terminals. While we did observe that SSp consistently drives MSNs, we did not observe evidence of strong connections between SSp terminals and CINs (**Suppl. Fig. 7**).

In summary, the *ex vivo* electrophysiology data functionally confirmed the anatomical findings from **Figure 3**. While MSNs received inputs from all areas tested, we observed a heavy bias in connectivity towards frontal cortical areas when recording from CINs.

### Frontal cortical areas, but not sensory, produce local acetylcholine-dependent dopamine release

Our results in **Figure 1** provide *in vivo* evidence that the visual stimulus-evoked dopamine responses depend on acetylcholine. Therefore, we next sought to determine whether the cortical inputs identified in **Figure 3** and tested functionally in **Figure 4** were capable of driving striatal dopamine release via an acetylcholine-dependent mechanism. We expressed an excitatory opsin (either ChR2 or ChrimsonR) in various cortical areas of C57B6/J mice (**Fig. 5a; Methods**) and used *ex vivo* fast scanning-cyclic voltammetry to measure dopamine transients elicited by optogenetic stimulations (470 nm, 5 ms, 1.3 mW; **Fig. 5b-c**) versus short electrical pulses (0.2 ms, 400 uA), that served as positive control. Optogenetic stimulation of ACA terminals consistently produced dopamine transients in DMS, although electrical evoked dopamine transients in this area were slightly larger, likely due to the additional signal from direct stimulation of dopamine axons (**Fig. 5d-f**; optogenetic: mean 0.39 [0.29 - 0.49 bootstrap 95% CI] μM, n = 9 slices from 3 animals; electrical: 1.43 [1.00 - 1.47] μM, n = 9 slices from 3 animals). Stimulating PL corticostriatal terminals produced similar results (**Fig. 5g-i**; optogenetic: 0.49 [0.39 - 0.56] μM, n = 10 slices from 4 animals; electrical: 0.79 [0.67 - 1.03] μM, n = 7 slices from 3 animals). To confirm that the cortically-evoked dopamine signal depended on CIN recruitment, we washed on a high concentration of DHβE (1 μM), a nicotinic acetylcholine receptor antagonist. Application of DHβE completely abolished any optogenetic evoked dopamine transients and significantly reduced electrical evoked events (**Fig. 5f**; ACA optogenetic DHβE: 0.011 [0.0096 - 0.025] μM, n = 6 slices from 3 animals; ACA electrical DHβE: 0.65 [0.42 - 0.86] μM, n = 6 slices from 3 animals; ACA optogenetic ACSF vs DHβE, p = 0.001; electrical ACSF vs DHβE, p = 0.007, Mann Whitney U test; **Fig. 5i**; PL optogenetic DHβE: 0.026 [0.0064 - 0.063] μM, n = 5 slices from 3 animals; PL electrical DHβE: 0.45 [0.27 - 0.62] μM, n = 5 slices from 3 animals; PL optogenetic ACSF vs DHβE, p = 0.0007; PL electrical ACSF vs DHβE, p = 0.02, Mann Whitney U test). In contrast, we found that optogenetic stimulation of sensory (VISp, or AUDp) corticostriatal terminals failed to produce any measurable dopamine transients in DMS (**Fig. 5j-l**, VISp: 0.015 [0.013 - 0.30] μM, n = 19 slices from 8 animals; **Fig. 5m-o**, AUDp: 0.011 [0.0098 - 0.054] μM, n = 19 slices from 4 animals). Interleaved electrical stimulation consistently produced dopamine transients in the same brain slices and under the same conditions, serving as positive controls and demonstrating feasibility (**Fig. 5k-l**, VISp: 0.67 [0.65 - 1.08] μM, n = 20 slices from 8 animals; **Fig. 5n-o**, AUDp: 0.33 [0.30 - 0.67] μM, n = 19 slices from 4 animals).

**Figure 5.**
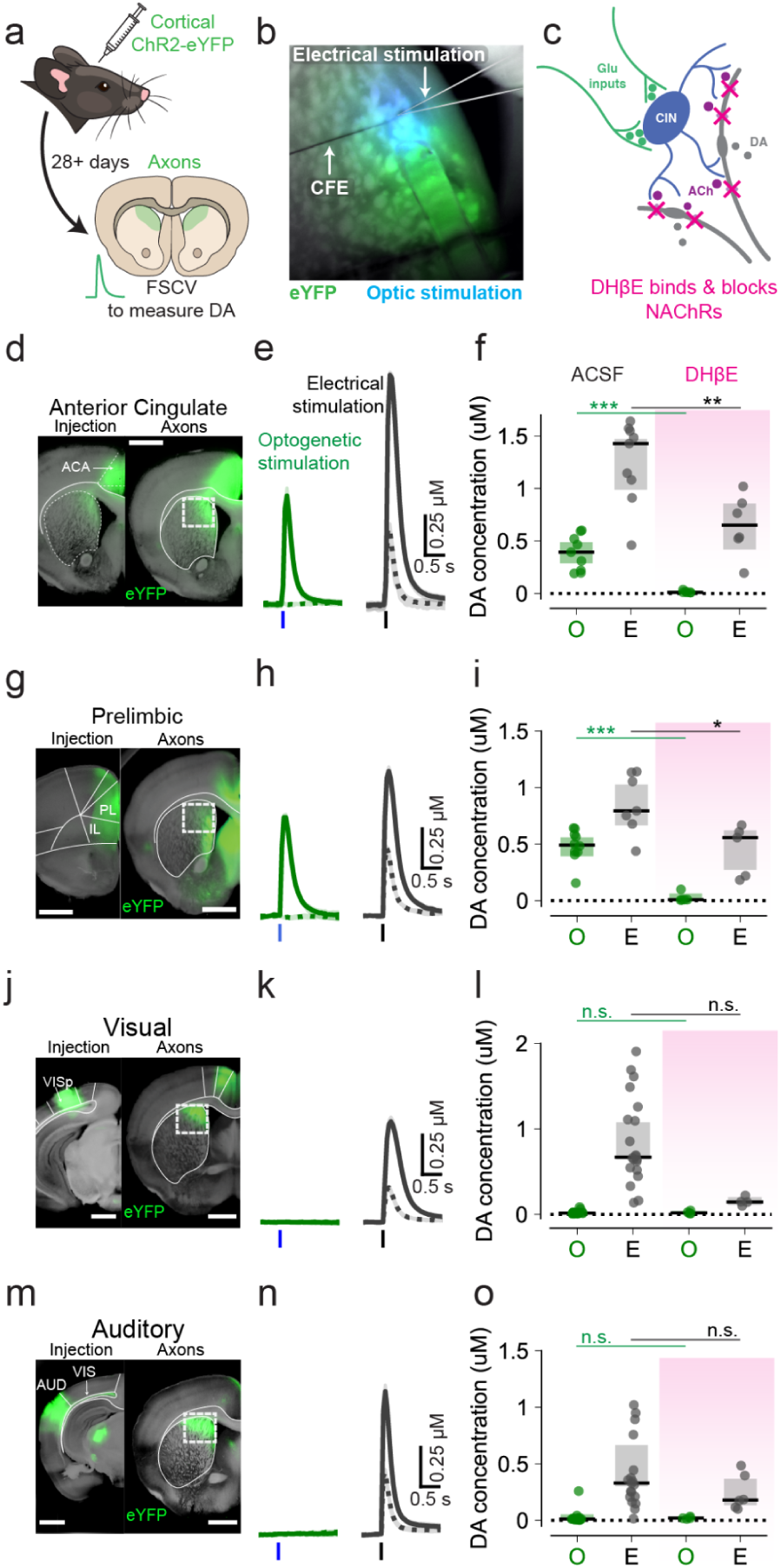
Corticostriatal projections from frontal, but not visual/auditory (sensory), cortical areas can evoke ex vivo local dopamine release in DMS. **a** Mice expressing ChR2-eYFP in frontal or sensory cortical areas were used for ex vivo brain slice recordings of dopamine with fast-scan cyclic voltammetry. **b.** Fluorescent image of brain slice shows ChR2-eYFP (green) expression in DMS from cortical projections and placement of carbon fiber electrode (CFE) used for fast-scan cyclic voltammetry to measure dopamine and glass pipette for electrical stimulation. Optogenetic stimulation was delivered via optic fiber (blue light). **c.** Diagram of the striatal microcircuitry proposed: cholinergic interneurons (blue) are recruited by glutamatergic corticostriatal inputs (gray) to release acetylcholine (purple), which activates nAChRs on DA axons (green) to evoke local dopamine release. DHβE application blocks nAChRs to prevent DA release. **d, g, j, m.** Left, ChR2-eYFP expression (green) in cortical injected area: anterior cingulate cortex (d), prelimbic cortex (g), visual cortex (VIS, j), auditory cortex (AUD, m). Right, ChR2-eYFP expressing projections (green) from the respective cortical areas in dorsomedial striatum. The white box shows the site of dopamine recordings. Scale bars are 1 mm. **e, h, k, n.** Single-slice average of dopamine transients measured with fast-scanning cyclic voltammetry (FSCV) and evoked by either optogenetic stimulation of corticostriatal projections (green) or electrical stimulation (gray) in the same recording site. Dashed lines show the remaining dopamine responses after washing in a nicotinic receptor blocker (1 μM DHβE). Traces are mean and shaded area are ± SEM). **f, i, l, o** Average amplitudes of dopamine transient in DMS elicited by electrical stimulation (gray bars) and optogenetic stimulation (green bars) before and after DHβE bath application (pink shade) for animals expressing ChR2-eYFP in ACA (f), PL (i), VIS (l,), and AUDp (o) cortical regions. Dots represent data from individual slices. Lines and shaded bars are median ± bootstrap 95% CI. *, denotes p < 0.05; **, denotes p < 0.01; ***, p < 0.001; non-significant pairs are not shown, Mann-Whitney U test.

Together, these experiments provide *ex vivo* evidence that, unlike sensory cortices, frontal cortical areas such as ACA and PL produce dopamine release through a local mechanism requiring nAChRs, and further suggest that these select corticostriatal inputs can drive synchronized activation of CINs.

### Corticostriatal projections from frontal cortices respond to visual stimuli and can recruit CINs and drive striatal dopamine and acetylcholine *in vivo*

We next returned to the *in vivo* preparation to ask whether PL/ACA axons projecting to DMS are activated in response to sensory stimuli (**Fig 6a-b**). We expressed GCaMP8s in one hemisphere of the prelimbic cortex and recorded calcium signals bilaterally from the DMS (**Fig 6 c-d**). The ipsilateral hemisphere to the injection responded strongly to the light flash (**Fig 6 e-f, left**). The contralateral hemisphere, which receives fewer projections than the ipsilateral hemisphere, responded more weakly to the light flash (**Fig 6e-f, right**).

**Figure 6.**
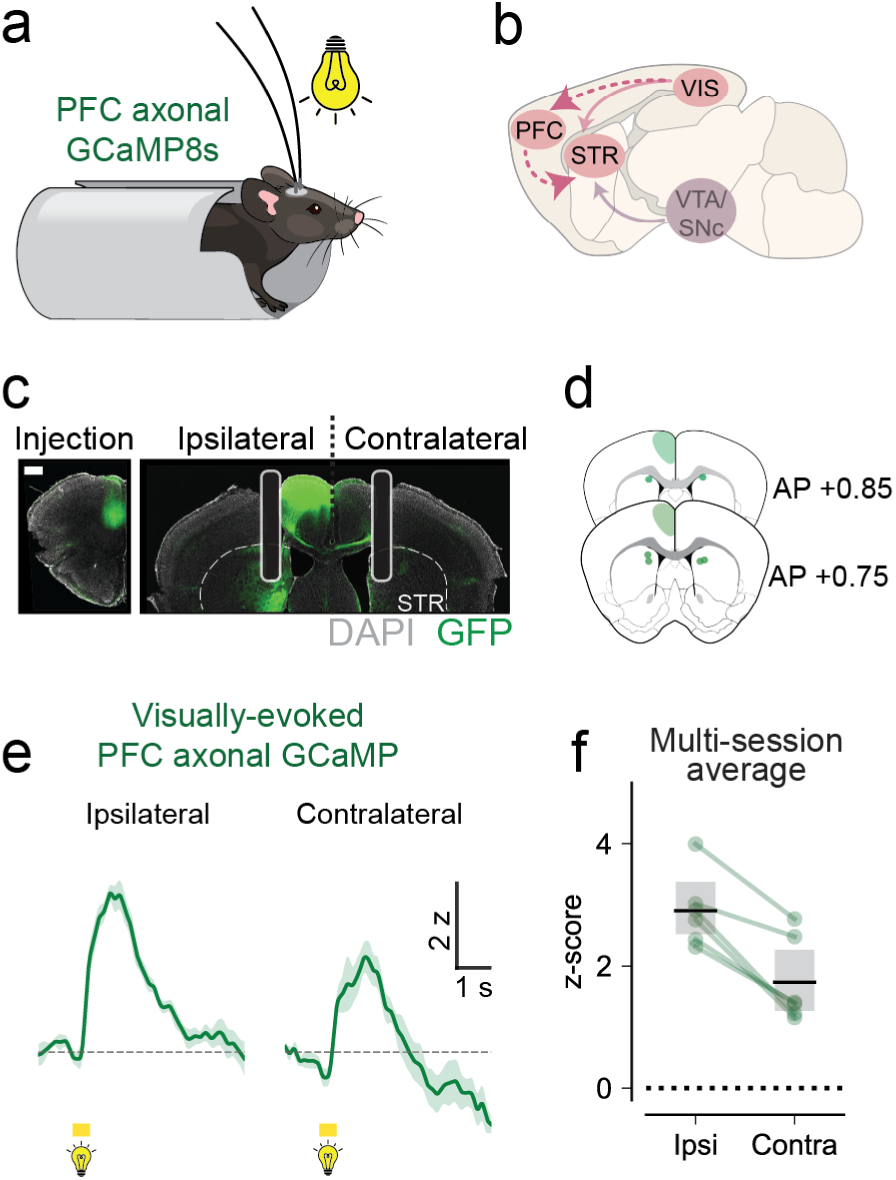
Visual stimuli produce time-locked activation of PFC corticostriatal projection. **a** GCaMP8s was expressed unilaterally in prelimbic cortex and recording fibers implanted bilaterally in DMS. Animals were headfixed and presented with 500ms visual stimuli, at least 6 weeks after surgery. **b.** Diagram depicts a canonical pathway (purple) from VTA/SNc for nicotinic-independent striatal dopamine in DMS following visual stimuli. A putative additional pathway (pink) from PFC for generation of nAChR-dependent striatal dopamine in DMS following visual stimuli. **c.** Representative images of unilateral GCaMP8 expression in prelimbic cortex (right, injection site) and green fluorescent projections into DMS showing bilateral fibers (right). **d,** Fiber placements in DMS for the animals used in the experiment. **e.** Representative average traces of visual-evoked calcium signals in PL axons in ipsilateral and contralateral DMS, relative to injection site. **j, l.** Average z-score amplitude of PL axon calcium responses timelocked to light flashes recorded from the ipsilateral hemisphere (j) and contralateral hemisphere relative to PL injection (l).

Given that frontal corticostriatal projections carry visual information, we next sought to test whether *in vivo* optogenetic stimulation of PL/ACA terminals in DMS could recruit CINs and evoke acetylcholine signals, as seen with visual stimulation. ChAT-IRES-cre mice were injected bilaterally in PL/ACA with ChrimsonR-tdTomato and in the DMS with either a cre-dependent calcium sensor (DIO-GCaMP8) or acetylcholine sensor (GRAB-ACh) in separate hemispheres (**Fig. 7 a-b**). These mice were implanted bilaterally with fiber-optic cannulae, through which we could simultaneously record fluorescent changes and stimulate cortical terminals in DMS with short pulses of red light (625 nm, 5.5 mW; single pulse or trains of 3 and 10 pulses at 20 Hz, 30s interval).

**Figure 7.**
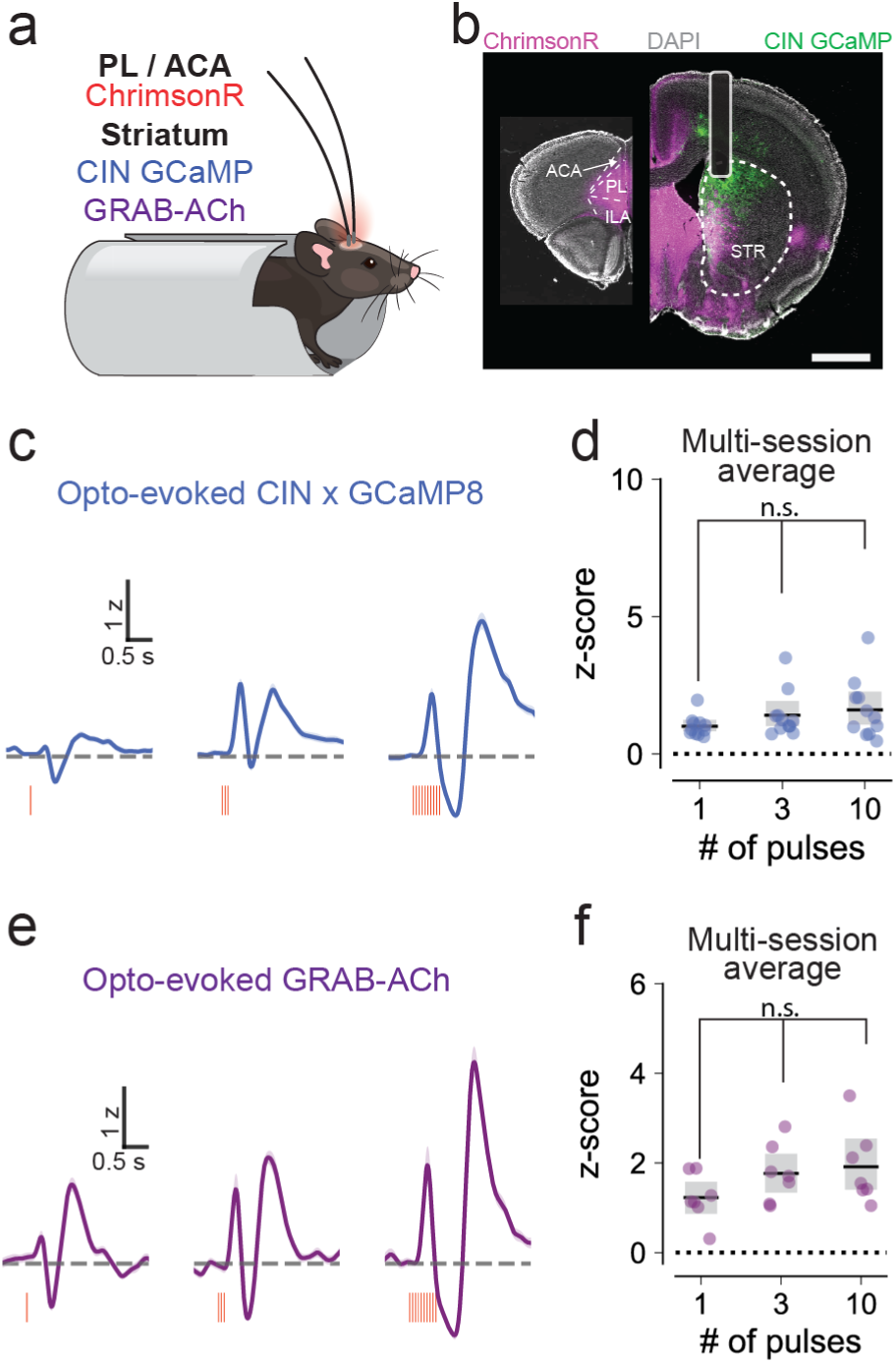
Corticostriatal projections from PFC recruit striatal CINs and drive acetylcholine release. **a.** Head-fixed set up used for experiments for optogenetic stimulation of PFC corticostriatal projection in ChAT-IRES-Cre mice expressing GCaMP8 in CINs and wildtype mice expressing GRAB_ACh in DMS. **b.** Example images of ChrimsonR-tdTomato expression in frontal cortical areas (pink) and their projection into dorsomedial striatum also showing GRAB_ACh expression (green). **c, e.** Representative traces of CIN GCaMP (c), and GRAB-ACh (e) responses to single pulse (10 ms,625 nm), trains of three pulses at 20 Hz (center), or trains of 10 pulses at 20 Hz (right). **d, f.** Maximum z-scored responses evoked by optogenetic stimulation of PFC projection in DMS for CIN GCaMP (d, n =11 mice) and GRAB_ACh (f, n = 7 mice. Dots represent data from individual mice. Lines represent the mean and gray bars represent the bootstrapped 95% confidence interval.. **f, i, k.** Maximum z-scored responses evoked from each animal expressing dLight1.3b (g), CIN GCaMP (i), and GRAB-ACh (k).

Stimulation of PL/ACA projections produced robust responses from both the calcium and acetylcholine sensors. These responses were also biphasic, consistent with the pattern seen in Figure 2 and possibly aligned with the characteristic burst-pause-burst pattern of striatal CINs (**Fig. 7c-f**). As with sensory cues, the later phases of these multiphasic responses were heterogeneous (**Fig. S8**). Therefore, for the quantitative analysis, we selected a time window corresponding to the first burst. A single pulse of PL/ACA terminal stimulation produced both CIN calcium responses (**Fig. 7c-d**; mean 1.0 [0.83 - 1.21 bootstrap 95% CI] z-score, n = 13 hemispheres from 11 animals) and acetylcholine responses (**Fig. 6e-f**; mean 1.23 [0.87 -1.59] z-score, n = 7 hemispheres from 7 animals). The amplitude of the first phase did not scale significantly with increasing stimulation (CIN GCaMP, 1 pulse: 1.00 [0.83 - 1.21] z-score; 3 pulses: 1.64 [1.12 - 2.41] z-score; 10 pulses: 1.84 [1.18 - 2.75] z-score, p = 0.09 one-way repeated measures ANOVA; GRAB-ACh, 1 pulse: 1.23 [0.86 - 1.58] z-score, 3 pulses: 1.77 [1.34 - 2.22] z-score; 10 pulses: 1.92 [1.40 - 2.53] z-score, p = 0.08 one-way repeated measures ANOVA). Consistent with the proposed local circuit mechanism, neither sensor response decreased significantly after receiving a systemic injection of a nAChR blocker or vehicle (**Suppl. Fig. 9**).

Lastly, we asked whether stimulation of prefrontal corticostriatal terminals produced cholinergic-dependent dopamine release. Wildtype mice were injected bilaterally with ChrimsonR-tdTomato in the PL/ACA and dopamine sensor, dLight1.3b, in the DMS with bilateral fiber implants and headframes (**Fig 8a-c**). A subset of these mice were also implanted unilaterally with opto-fluid cannula to allow for intracranial delivery of mecamylamine during recordings (**Fig 8d**). All mice received the same optogenetic stimulation protocol described above. Even a single 10-ms pulse of red light evoked measurable striatal dopamine responses (**Fig. 8e-f**, mean 3.02 2.95 - 3.75 bootstrap 95% CI] z-score, n = 19 animals, averaged across sessions). The amplitude of the dopamine response increased as the number of pulses increased, with evidence of saturation and prolonged duration with the longest stimulation train **(Fig. 6e-f**, 3 pulses: 5.65 [5.34 - 7.07] z-score; 10 pulses: 6.88 [6.45 - 8.75] z-score, averaged across sessions; p < 0.0001 one-way repeated measures ANOVA). We repeated these experiments in DLS with stimulation of SSp terminals and observed that stimulation produced small responses in CIN GCaMP and GRAB-ACh, but was unable to evoke positive dLight responses (**Suppl. Fig. 10)**.

**Figure 8:**
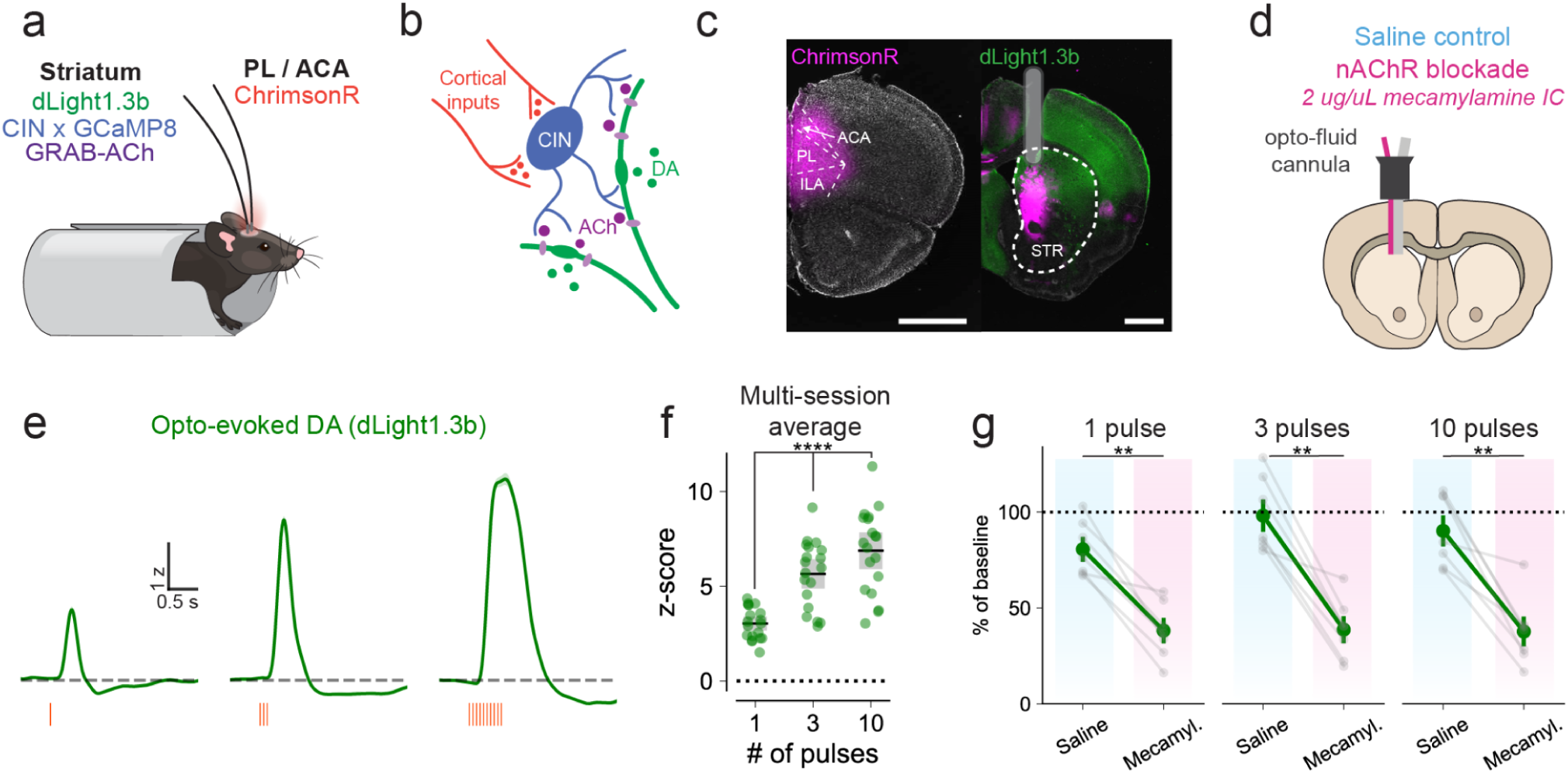
Stimulation of PFC corticostriatal projections triggers nAChR-dependent dopamine release in DMS. **a** Head-fixed experimental setup used for optogenetic stimulation of corticostriatal projections expressing Chrimson-tdTomato in prelimbic or anterior cingulate cortex, bilaterally. Fiber implants in DMS bilaterally allow for optogenetic stimulation and recording of dLight, or GCaMP8 expressed in CINs, or GRAB-ACh expressed in the dorsomedial striatum. **b.** Diagram of the proposed striatal microcircuitry: striatal cholinergic interneurons (blue) are recruited by glutamatergic inputs from corticostriatal ChrimsonR-expressing projections (orange) to release acetylcholine (purple), which activates nAChRs on DA axons (green) to produce local dopamine release. **c.** Example images of ChrimsonR-tdTomato expression in frontal cortical areas (magenta) and their projection into dorsomedial striatum where dLight1.3b (green) was also expressed. **d.** A subset of dLight-expressing mice were implanted with optofluid cannula to allow intracranial infusions of saline or mecamylamine in DMS. **e.** Representative traces of dLight1.3b responses evoked by optogenetic stimulation of PFC projections with a single pulse (10 ms, 625 nm) (left) and by trains of three pulses at 20 Hz (center) or trains of 10 pulses at 20 Hz (right). **f.** Maximum z-scored dLight1.3b responses evoked by optogenetic stimulation of PFC projection in DMS (n= 11 mice). Green dots represent data from individual hemispheres. Lines represent the mean and gray bars represent the bootstrapped 95% confidence interval. **g.** Effect of intracranial infusion of saline (blue) or mecamylamine (pink) on optogenetically-evoked dLight responses following single pulse, trains of 3 and 10 pulses stimulation. Amplitudes were normalized to each animal pre-infusion baseline (n= x hemispheres/y mice).

To test whether a component of this optogenetic evoked response was dependent on acetylcholine, a subset of mice received intracranial infusions of mecamylamine at the recording site (2 ug/uL at a rate of 0.06 uL/min). Blocking nAChRs produced significant decreases in dopamine at all stimulation levels (**Fig. 8g**; single pulse reduced to 38% ± 5.6%, 3 pulses reduced to 39% ± 5.9%, 10 pulses reduced to 38% ± 6.7%; treatment p = 0.0003, stimulation intensity p = 0.04, interaction p = 0.15, two-way repeated-measures ANOVA). Whereas only some mice received intracranial infusions, all mice received systemic intraperitoneal injections of 10 mg/kg mecamylamine (**Suppl. Fig. 9**). Systemic mecamylamine was not effective at suppressing the optogenetic-evoked dopamine (0% suppression vs 62% suppression with intracranial), unlike with visually-evoked dopamine responses. This may be due to a combination of factors: (A) the stronger over-drive of the projections achieved with optogenetic stimulation, and/or (B) the lower effective dose of blocker achieved by systemic administration, capped by peripheral effects. Altogether, these *in vivo* experiments offer compelling evidence that brief activation of PL/ACA projections to the dorsomedial striatum can evoke cholinergic-dependent dopamine signals, via the local recruitment of CINs and subsequent ACh release.

## Discussion

This study provides new insights into how salient sensory stimuli control dopamine transmission in the striatum. Using a combination of anatomical and functional experiments, both *in vivo* and *ex vivo*, we identified which corticostriatal inputs can modulate striatal dopamine, and which cannot. We show that visual stimuli elicit striatal dopamine release in part through a cholinergic-dependent mechanism involving recruitment of CINs and acetylcholine release in the dorsomedial striatum. Despite the presence of prominent projections from the primary visual and auditory cortices to the dorsomedial striatum, these cortical areas do not connect strongly to striatal CINs and are unable to drive dopamine release. In contrast, frontal cortical areas such as the prelimbic and anterior cingulate cortices robustly recruit striatal CINs and acetylcholine to drive dopamine release *ex vivo* and *in vivo*. In mice, studies have demonstrated that the medial frontal cortex is increasingly activated by visual events throughout learning, and that inactivating this area dramatically affects an animal’s ability to perform visual tasks^46–48^. Thus, the current study identifies a key distinction between primary sensory and frontal cortical inputs to the striatum, with important implications for how the brain processes salient stimuli that support associative and reinforcement learning.

Previous studies showed that certain cortical and thalamic inputs can evoke striatal dopamine *ex vivo*, but two key questions remained: Which cortical areas selectively recruit CINs to regulate dopamine? Does this mechanism operate *in vivo* under behaviorally relevant conditions? Slice experiments demonstrated that inputs from the motor cortex and parafascicular thalamus strongly engaged CINs in the dorsal striatum and triggered dopamine-dependent plasticity^19^. The retrograde monosynaptic rabies tracing from our study supports these findings and identifies the dorsolateral and dorsocentral striatum as the target of the motor cortex and parafascicular thalamus inputs to CINs. Subsequent studies extended the findings to prelimbic cortex and intralaminar thalamus inputs^20^; and again our anatomical and functional experiments here confirm those findings and further identify the dorsomedial subregion as the target from these PL/ACA inputs to CINs. *In vivo* stimulation of the prelimbic cortex was also shown to increase striatal glutamate, acetylcholine, and dopamine using microdialysis^20^, though with insufficient temporal precision to link dopamine release to behavioral events. Our results here address this gap by providing *in vivo* evidence of frontal cortex (PL/ACA) control over striatal dopamine with higher temporal precision. Crucially, we further demonstrate the ability of these corticostriatal inputs to recruit CINs and drive dopamine via nicotinic-dependent mechanisms. Functional and anatomical connectivity to CINs is restricted to specific cortical (frontal and motor) and thalamic inputs and we show that it is absent in primary sensory cortices.

It is important to mention that *in vivo* optogenetic stimulation of cortical axons in the striatum is likely to also activate passing axons and axon collaterals that synapse onto other targeted areas. More specifically, PL/ACA projections to the dorsomedial striatum also send collaterals to dopaminergic midbrain regions, which likely contribute to a portion of the striatal dopamine signals we measured *in vivo*^49^. Also note that a recent study showed muscarinic receptors contribute to cholinergic regulation of dopamine in vivo, which could be responsible for the portion of the optogenetic response that remains after infusing mecamylamine (a nicotinic acetylcholine receptor antagonist)^50^. Thus, while not acting alone, a number of our experiments provide evidence for a local mechanism dependent on CINs and acetylcholine. First, intracranial local infusions of nAChR blocker mecamylamine successfully reduced dopamine responses evoked by both cue- and optogenetic stimulation of PL corticostriatal projections. Systemic mecamylamine dramatically reduced cue responses, but only slightly reduced the optogenetic-evoked dopamine responses, perhaps because the dose limitation for systemic administration was more easily overcome by the optogenetic over-stimulation of axons. Second, visual stimuli and ACA/PL projections also recruit CIN and drive acetylcholine release in vivo, as expected for local cholinergic-dependent dopamine release. Third, our *ex vivo* experiments isolated the local striatal microcircuitry and confirmed that corticostriatal projections from PL/ACA form strong functional synapses onto CINs and robustly recruit them to evoke local dopamine release, consistent with a direct cortical effect within the striatum. Because all axons from neurons projecting to the striatum are severed in *ex vivo* brain slice preparations, only the axonal projections within the slice are activated during the experiments. Any direct projections from cortex to the midbrain dopamine neurons were excluded and cannot be responsible for the dopamine signals measured *ex vivo*. A last caveat to consider is that the acetylcholine release in the slices may arise from extra-striatal sources of acetylcholine axons, rather than striatal CINs. However, other studies have found that projections from pedunculopontine nucleus or laterodorsal tegmentum do not produce dopamine signals in the striatum^51^. Altogether, we conclude that the unique ability of PL/ACA to evoke dopamine likely arises from the strength of its connections with CINs, a conclusion supported by both anatomical and functional data.

A strength of this study is its integration of dopamine, calcium, and acetylcholine measurements. However, direct comparison of signal magnitude and timing across these sensors is difficult because each differs in affinity and kinetics. At present, we can only interpret these timing differences cautiously. Importantly, a recent study examined the temporal relationship between acetylcholine and dopamine in vivo using precisely timed optogenetic stimulation of CINs combined with ultrafast dopamine imaging and identified a biphasic dopamine response^50^. The initial dopamine peak depended on β2-containing nicotinic receptors, which are high-affinity receptors expected to respond to even small increases in acetylcholine. Together, these findings support the idea that relatively small increases in acetylcholine may be sufficient to evoke dopamine release, as observed with single-pulse stimulation.

In considering our anatomical tracing data, it is important to note that rabies tracing results varied depending on the striatal location of the starter cells. When we targeted the anterior dorsomedial striatum, we observed projections from PL/AC to CINs. When we targeted central or lateral portions of the anterior striatum, we observed projection to CINs from the motor cortex and some somatosensory cortical areas. Thus, overall there was a parallel pattern of medial-to-lateral segregation of inputs to CINs. Our tracing data refine previous work by detailing the cortical and thalamic sources of input to CINs within specific striatal subregions using an improved ChAT-IRES-Cre mouse line without common events of spontaneous recombination^54^. Different patterns would be expected for starter cells in the rostral portion of striatum, the tail, or the nucleus accumbens. Future studies comparing these subregions are needed to assess whether similar connectivity rules apply.

While rabies tracing identifies the anatomical distribution and laminar origin of inputs to CINs, it cannot quantify the functional efficacy of the connections. To assess the functional synaptic strength, we therefore used electrophysiology. Our whole-cell recordings showed that CINs receive rare and weak inputs from visual and barrel cortices, whereas neighboring MSNs receive reliable synapses. Nonetheless, it remains possible that coincident activation from multiple sensory areas could summate to recruit CINs and evoke dopamine release under some specific conditions. Broadly, our functional results aligned with the anatomical data: rabies tracing revealed that CINs in the dorsomedial striatum (DMS) receive dense input from frontal regions, particularly ACA, whereas CINs in the dorsolateral striatum (DLS) are more strongly innervated by motor and somatosensory cortices. This topographical organization parallels the connectivity of cortical inputs to striatal MSNs^52–54^, reinforcing the view that the DMS is engaged in cognition and goal-directed behavior^55,56^, while the DLS supports action selection and habits^3,57–59^.

Our results provide direct support for the idea that visual stimuli and cortical inputs to the striatum can evoke dopamine release through a local circuit involving CINs. We acknowledge that the relationship between dopamine and acetylcholine is complex and remains controversial. Two studies showed that striatal acetylcholine and dopamine signals fluctuate in an anti-correlated manner *in vivo*^36,37^. These observations were further supported by recent findings from *ex vivo* slice experiments showing acetylcholine may, in some contexts, inhibit midbrain-evoked dopamine^21^, which could explain the anticorrelation observed in the *in vivo* signals. One of these studies reported that intracranial infusions of DHβE, a nicotinic receptor blocker, did not affect spontaneous dopamine signals, nor reward-driven signals^36^, which midbrain neurons would be expected to drive. Meanwhile, inhibiting acetylcholine release from CINs, striatum-wide, had significant effects on behavior and behaviorally evoked dopamine signals^37^. However, more local inhibition of acetylcholine release in the ventrolateral striatum had no effect in the same contexts^37^. These studies were also performed in dorsolateral striatum and central striatum. Our results provide evidence for a local CIN-dependent circuit mechanism specifically within the DMS, and additional efforts will be required to understand possible regional differences in striatal circuit mechanisms that may eventually settle this important controversy.

More recent work in the nucleus accumbens shows compelling evidence that the local microcircuitry regulating striatal dopamine signals plays a role during both effortful tasks^35^ and active avoidance behavior^60^. These studies align well with the current findings from our study in which these local acetylcholine-dependent dopamine signals may work together and summate with midbrain-originated dopamine signals. Understanding the temporal dynamics of this cholinergic-dependent dopamine signal and how it integrates with midbrain-evoked dopamine is an important next step. Also, a deeper understanding of how CIN-dependent dopamine release contributes to the dopamine tone *in vivo* may clarify the long-standing discrepancies between tone and phasic dopamine regulation and functional role. Lastly, we highlight a need to further explore the role of this acetylcholine-mediated control over dopamine throughout learning, stimulus-reward associations and also reinforcement learning, which may account for the historic role of CIN firing pattern in cue-reward learning.

In summary, our study reveals that sensory stimuli and frontal cortical regions, including the prelimbic and anterior cingulate cortices, robustly engage CINs and elicit acetylcholine signals to trigger striatal dopamine release in the anterior dorsomedial striatum. Primary sensory cortices, while projecting to the same striatal subregion, do not recruit CINs nor share the ability to drive striatal dopamine. Thus, we reveal a key distinction between sensory and frontal cortical inputs, identifying the latter as privileged drivers of striatal dopamine that may support associative and reinforcement learning. Disentangling the mechanisms behind cholinergic control over striatal dopamine and associated learning could prove critical to understanding the root cause of disorders involving dysregulation of striatal dopamine.

## Author Contributions

Conceptualization, H.C.G., V.A.A., and R.J.K.; Methodology, H.C.G., C.R.G., R.J.K., and V.A.A.; Formal Analysis and Data Curation, H.C.G. and C.R.G; Investigation, H.C.G., R.R., E.S.S., J.H.S., M.E.A., L.G.A., H.B.K., L.M.A., R.P., and C.R.G.; Resources, V.A.A, R.J.K., and C.R.G.; Writing - Original Draft, H.C.G.; Visualization and Writing - Review & Editing, H.C.G., R.J.K., and V.A.A.; Supervision, V.A.A. and R.J.K.; Funding Acquisition, V.A.A., R.J.K., C.R.G, and H.C.G..

## Competing Interest Statement

The authors declare no competing interests.

## Materials and Methods

### Animals

All experimental procedures were approved by the NIH Institutional Animal Care and Use Committee (IACUC) and complied with Public Health Service policy on the humane care and use of laboratory animals. Both female and male mice were used for all experiments. Electrophysiology experiments used either a ChAT-IRES-Cre (Δneo) line^61^ (for CIN or MSN recordings, B6.129S-*Chat^tm1(cre)Lowl^*/MwarJ; RRID: IMSR_JAX:031661), a Pvalb-IRES-Cre line (for MSN recordings, B6.129P2-Pvalbtm1(cre)Arbr/J; RRID: IMSR_JAX:017320), or a SST-IRES-CRE line^62^ (for MSN recordings, B6J.Cg-Ssttm2.1(cre)Zjh/MwarJ; RRID: IMSR_JAX:028864) crossed with an Ai14 Cre-dependent tdTomato line^63^ (B6.Cg-*Gt(ROSA)26Sor^tm14(CAG-tdTomato)Hze^*/J; RRID: IMSR_JAX:007914). Fast-scanning cyclic voltammetry experiments used wild-type C57B6/J mice or the ChAT-IRES-Cre (Δneo) x Ai14 cross. Rabies experiments were done in the ChAT-IRES-Cre (Δneo) line. C57BL6/J, DAT-IRES-cre^64^ (B6.SJL-Slc6a3tm1.1(cre)Bkmn/J, RRID: IMSR_JAX:006660), ChAT-IRES-Cre (Δneo), pDyn-IRES-cre^65^ (B6;129S-Pdyntm1.1(cre)Mjkr/LowlJ, RRID:IMSR_JAX:027958), and Drd1a-Cre (B6.FVB(Cg)-Tg(Drd1-cre)EY262Gsat/Mmucd, RRID: MMRRC_030989-UCD) mice were used for *in vivo* fiber photometry experiments.

### Fiber photometry

#### Surgeries

Animals (N = 63, 41 males, 22 females; age: 3 - 9 months, average 5 months, **Table S1**) were injected with AAV5-hSyn-dLight1.2^66^ (titer = 2.3E+13; Addgene #111068-AAV5; RRID:Addgene_111068), AAV9-hSyn-dLight1.3b^66^ (titer = 2.5E+13; Addgene #135762-AAV9; RRID:Addgene_135762), AAV9-Syn-FLEX-jGCaMP8s-WPRE^67^ (titer = 2.7E+13, Addgene #162377-AAV9; RRID: Addgene_162377), AAV9-Syn-FLEX-jGCaMP8f-WPRE^67^ (titer = 2.0E+13, Addgene #162379-AAV9, RRID: Addgene_162379) and/or AAV9-Syn-GRAB-ACh3.0(4.3)^68^ (titer =6.7E+12, WZ Bioscience #YL001003-AV9-PUB) in DMS (400-500 nL at 180 nL / min; +0.9 mm AP, +/- 1.3 mm ML, -2.8 mm DV from Bregma). Mice used for midbrain dopamine cell body recordings were injected with AAV9-Syn-FLEX-jGCaMP8s-WPRE in the VTA / SN (500 nL at 180 nL / mine; -3.3 mm AP, - 0.6mm ML, - 4.5 mm DV). A subset of these mice were additionally injected with AAV5-hSyn-ChrimsonR-tdTomato^69^ (titer = 1.20E+13; Addgene #59171-AAV5; RRID:Addgene_59171) in prelimbic cortex (120 nL at 120 nL / min; +2.1 mm AP, +/- 0.4 mm ML, -2.3 mm DV from Bregma) or anterior cingulate cortex (120 nL at 120 nL / min; +1.0 AP, +/- 0.3 mm ML, -1.5 mm DV from Bregma). A custom headpost and optic fiber cannulae (DMS: 400um Ø, 0.37 NA, 2.5 mm long; VTA/SN: 200um Ø, 0.37 NA, 4.5 mm long; Doric) were implanted above the injection sites using Metabond (C&B Parkell).

#### Photometry system

Thorlabs LED drivers (LEDD1B) were used to drive Thorlabs 405 nm (M405F1), 470 nm (M470F4), and 625 nm (M625F2) fiber-coupled LEDs connected to a six-channel Doric filter cube (FMC6_IE(400-410)_E1(460-490)_F1(500-540)_E2(555-570)_F2(580-680)_S) by 0.5 NA / 400 μm Ø patch cords (Thorlabs, M301L01). Subject cords were custom-ordered from Doric (MFP_400/440/1100-0.37_1m_FCM-ZF1.25(FP)_LAF). Newport Femtowatt Receivers (Doric, NPM_2151_FOA_FC) were connected to the filter cubes using custom 0.5 NA / 600 μm Ø patch cords (Thorlabs). Femtowatt receivers fed into a Tucker Davis Technologies RZ5P processor. All fiber optic cords were photobleached for 24 hours before the start of the experiment. 405 nm and 470 nm LEDs were driven at 215 and 326 Hz. Data were collected at 1017 Hz and demodulated using Tucker Davis Technologies Synapse software. An Arduino interfaced with the 625 nm LEDs and the RZ5P processor controlled and timestamped optical stimulation.

#### Sensory and optogenetic stimulation

405 nm and 470 nm LEDs were calibrated daily to 20 μW and 30 μW, respectively, as measured at the fiber tip. Mice were head-fixed in an acrylic tube for the duration of recordings. Mice were headfixed into a custom head fixation setup with their body resting within an acrylic tube. All behavioral tasks were performed in a dark sound-attenuated behavior chamber (Med Associates) fitted with 20mm black sound-insulating foam. All behavioral tasks and external stimuli were controlled using pyControl^70^ (Open Ephys Production Site). Mice only experienced each stimulus five times per session with a maximum of two sessions per day, to avoid habituation.

Auditory cues were delivered at 7.5 KHz at 70 dB for 500 ms (a volume level comparable to normal conversation). Tones were delivered via an 8 ohm speaker (DigiKey; part number GF0401M-ND) placed 6 inches directly above the mouse’s head. The speaker was connected to a microcontroller (ATmega328P) where tones were generated using Arduino’s tone function. Visual cues were delivered via a pyControl houselight LED that uses three 16 lux LEDs on a plastic strip, placed in the behavior chamber above / slightly in front of the mouse. The luminance of the LED was measured to be 5.9 cd/m^2^ on the floor of the box, and 0.6 cd/m^2^ on the black foam sidewalls (light intensity comparable to a nightlight or natural light at dawn / dusk). The visual stimulus was intentionally dim to avoid a startle response that could be produced by a brighter light in contrast to the dark behavior box.

Optogenetic stimulation used 625 nm LEDs calibrated to 5.5 mW output. Optogenetic stimulation was delivered every 30 seconds in either single 10 ms pulses, 10 ms three-pulse trains at 20 Hz, or 10 ms ten-pulse trains at 20 Hz.

#### Pharmacology

Mecamylamine (2843/10, Tocris) was prepared fresh in 0.9% sterile saline and filtered for a final concentration of 2 ug / uL for intracranial infusion or 10 mg / 10 mL for intraperitoneal injection, allowing mice to be injected with a 10 mL / kg dosage. During pharmacology sessions, 7.5 minutes of optogenetic data were collected followed by 5 repetitions of each cue. For intracranial infusions, a SP101l syringe pump (world precision instruments) loaded with a 1700 series 10uL gas-tight syringe (Hamilton, 80008) was used to start the infusion (mecamylamine or saline vehicle) at a rate of 0.06 uL / min and allowed to flow for ∼10 minutes before the optogenetic stimulation and cue protocols were repeated. For intraperitoneal injections, mice were injected intraperitoneally with mecamylamine (10 mg / kg prepared for a final 10 mL / kg volume) or saline (10 mL / kg) and returned to their home cage for 30 minutes before repeating the optogenetic stimulation and sensory cue protocols.

#### Analysis

Data were converted to ΔF/F signals using GuPPy, an open-source Python package^71^ (RRID: SCR_02235345). ΔF/F was calculated by subtracting the isosbestic channel from the signal channel and dividing by the isosbestic. The same Python package was used to baseline-correct the traces and create PSTHs. The preprocessed data were then further analyzed and plotted in Python. Data were z-scored either over a single file (baseline non–pharmocological sessions) or over the “pre” and “post” files together (pharmacological sessions) by calculating the mean and standard deviation of the trace(s), subtracting the mean and dividing by the standard deviation. Baseline data were averaged per mouse across sessions. Pharmacological data were always collected and reported from a single session.

Peak amplitude of the evoked responses were calculated by finding the maximum value within 2 s from stimulation for dLight (which produced larger, longer responses with a single peak) or within a 0.6 s window from stimulation for GCaMP or GRAB-ACh (which tended to produce multiphasic responses with faster phases). Because later phases of the responses were heterogenous, we limited the analysis to the first phase of the response, which was consistent across animals.

#### Histology

Mice were deeply anesthetized with isoflurane and transcardially perfused with phosphate-buffered saline, followed by 4% paraformaldehyde (PFA). Brains were extracted, post-fixed in 4% PFA at 4 °C overnight, and transferred to 30% sucrose in PBS for >48 h at 4 °C. Brains were flash frozen in isopentane and stored at −80 °C. 35-µm coronal cryo-sections were cut using a cryostat and collected in PBS containing 0.01% sodium azide. To verify viral expression and fiber placement, brain sections were gently rocked for 3 × 10 min in PBS, blocked with 10% normal goat serum (NGS; Jackson Immuno, 005-000-121; RRID: AB_2336990) in PBS containing 0.1% Triton X-100 (PBS-Tx) for 1 h at room temperature, and then incubated in rabbit anti-GFP, 1:500 (Invitrogen, G10362; RRID: AB_2536526) primary antibody in PBS-Tx at 4 °C for 24 h. Sections were rinsed 3 × 10 min with PBS and incubated in 488 anti-rabbit,1:800 (Jackson Immuno, 111-545-003; RRID: AB_2338046) secondary antibody in PBS-Tx for 2 h at room temperature. Sections were washed 3 × 10 min with PBS, mounted on slides with coverslip and ProLong™ Gold antifade reagent with DAPI (Invitrogen, P36935). All slides were imaged using consistent exposure settings on a ZEISS Axioscan 7 slide scanner or a Keyence BZ-X810 microscope.

### Slice physiology

#### Surgeries

Animals (n = 47; 21 males, 26 females; age: 2-13 months, median 3 months) were injected with AAV5-hSyn-ChR2(H134R)-eYFP (titer = 1.80E+13; Addgene, #26973; RRID:Addgene_26973) and/or AAV5-hSyn-ChrimsonR-tdTomato (as above). PL injections (100-120 nL at 120nL/min) were targeted to +2.1 mm AP, +/- 0.4 mm ML, -2.3mm DV from Bregma. ACA injections (100-120 nL at 120nL/min) were targeted to +1.0 mm AP, +/- 0.3 mm ML, -1.5mm DV from Bregma. Somatosensory cortex injections (120 nL at 120nL/min) were targeted to -1.25 mm AP, +/- 3.0 mm ML from Bregma, and -0.5 mm DV from Pia. Visual cortex injections (120 nL at 120nL/min) were targeted to -3.5 mm AP, +/- 2.3 ML from Bregma, and -0.5 mm from Pia. Auditory cortex injections (120 nL at 120nL/min) were targeted to -2.7 mm AP, +/- 4.3 mm from Bregma, and -0.7 mm from Pia.

#### Slice preparation

Mice were anesthetized and decapitated at least four weeks post-surgery. Brains were extracted, mounted on a vibratome (VT-1200S, Leica Microsystems), and sliced coronally (240 μm) in oxygenated cutting solution heated to 32°C, containing the following (in mM): 90 sucrose, 80 NaCl, 24 NaHCO3, 1.25 NaH2PO4, 10 glucose, 3.5 KCl, 0.5 CaCl2, 4.5 MgCl2, and 3 kynurenic acid. Slices were incubated for 20 min at 32°C in ACSF containing the following (in mM): 124 NaCl, 1 NaH2PO4, 2.5 KCl, 1.3 MgCl2, 2.5 CaCl2, 20 glucose, 26.2 NaHCO3, and 0.4 ascorbic acid, and kept at room temperature after that until use. The recording chamber was perfused at 2 ml/min with ACSF heated to 32°C using an inline heater (Harvard Apparatus).

#### Fast scanning-cyclic voltammetry

FSCV was performed in the dorsal striatum. Carbon-fiber electrodes (CFEs) were prepared with a cylindrical carbon fiber (7 μm diameter, ∼150 μm of exposed fiber) inserted into a glass pipette. Before use, the CFEs were conditioned with an 8-ms-long triangular voltage ramp (−0.4 to 1.2 and back to −0.4 V vs Ag/AgCl reference at 400 V/s) delivered every 15 ms. CFEs showing current >1.8 µA or <1.0 µA in response to the voltage ramp at ∼0.6 V were discarded. CFEs were held at −0.4 V versus Ag/AgCl, and the same triangular voltage ramp was delivered every 100 ms. Using the same CFE and location, dopamine signals were evoked by alternating electrical and optical stimulation, delivered every 2 mins. For electrical stimulation, a glass pipette filled with ACSF was placed near the tip of the carbon fiber and a rectangular pulse (0.2 ms, 400 μA). For optogenetic stimulation, 470 nm blue light (1.3 mW) was delivered through either a 40x objective via a CoolLED (pE-800) or a fiber-optic patch cord (200 μm diameter, 0.22 NA, ThorLabs) attached to a ThorLabs fiber-coupled LED (M470F4, driven by a LEDD1B driver). The light source was placed over the CFE to deliver square light pulses (5-10ms). Data were collected with a retrofit headstage (CB-7B/EC with 5 mΩ resistor) using a Multiclamp 700B amplifier (Molecular Devices) after a low-pass filter at 3 kHz and digitized at 100 kHz using a data-acquisition board (NI USB-6229 BNC, National Instruments). Data acquisition and pre-processing were performed using custom-written software, VIGOR, in Igor Pro (Wavemetrics) using mafPC (courtesy of MA Xu-Friedman). Further analysis was conducted using Python. The current peak amplitudes of the evoked dopamine transients were converted to dopamine concentration according to a post-experimental calibration using 1-3 μm DA.

#### Electrophysiology

Striatal CINs were identified by fluorescence and confirmed by their characteristic spontaneous firing pattern. Whole-cell recordings were performed from CINs in the striatum using glass pipette electrodes with a resistance of ∼3-4 MΩ, filled with an internal solution (pH 7.25, 290-310 mOsm) containing the following (in mM): 120 CsMS, 10 CsCl, 10 HEPES, 0.2 EGTA, 10 sodium phosphocreatine, 4 Na2-ATP, and 0.4 Na-GTP. For input-output curves, internal solution also contained 4.4 mM lidocaine to block sodium channels and prevent firing. Cholinergic interneurons were held at a holding potential of -70 mV to keep these spontaneously firing neurons far away from firing threshold. Medium spiny neurons were held at a holding potential of -70 mV or -55 mV, where the cell input resistance is higher. All recordings were done in the presence of 5 μM CPP (Tocris, #0247) to block NMDA currents. Excitatory post-synaptic currents were recorded in response to 1 or 10 ms duration square light pulses (titrated based on synapse strength to avoid inducing action potentials; 10 ms was used when there was no response, to demonstrate that even very intense stimulations could not produce a response). A dual-color LED (Doric LEDC2_405/ 595) was used to alternate between pulses of violet-shifted (405 nm, 1.6 mW) or orange light (595 nm, 1.0 mW). Data were collected using a Multiclamp 700B amplifier after a low-pass filter at 1 kHz and digitized at 5 kHz using pClamp10 software (Molecular Devices).

#### Analysis

Data were analyzed in Python using pyABF (https://github.com/swharden/pyABF). Sweeps were filtered with a 100Hz 4th-order lowpass Butterworth filter to remove high-frequency noise. Data were baseline corrected to a standard window prior to stimulation, and noise was subtracted to the height of the average peak within that window. Peaks were detected as the minimum (EPSCs) within a 300 ms window from stimulation. For visualization, the absolute values of the EPSCs were plotted as positive numbers. All data are presented as the mean and SEM.

### Anatomical tracing

ChAT-IRES-Cre (Δneo) (n = 12, 7 males, 5 females; age: 2-21 months, median 2 months) were injected unilaterally in DMS or DLS, as described above, with helper viral vector AAV8-DIO-Ef1a-TVA-Flag-2A-N2cG (Addgene #172360-AAV8; RRID:Addgene_172360), and rabies retrograde vector EnVA-CVS-N2c-tdTomato-FlpO one week apart^72^. Six days after the second injection, mice were deeply anesthetized with isoflurane and transcardially perfused with phosphate-buffered saline, followed by 4% paraformaldehyde (PFA). Brains were removed, fixed overnight in 4% PFA and transferred to 20% sucrose in PBS. After the brains had equilibrated in 20% sucrose/PBS coronal sections were cut at 50 μm on a freezing microtome and collected in PBS. TdTomato labeling in trans-synaptically rabies labeled neurons was amplified immunohistochemically using sequential incubation in primary antibodies, rabbit anti-RFP (Rockland, 600-401-379) followed by secondary antibodies goat anti-rabbit Alexa Fluor 555 (ThermoFisher, A32732). Sections were mounted onto slides and labeled with the fluorescent blue Nissl stain Neurotrace 435 (ThermoFisher, N21479). All sections through the brain from the frontal pole to the brainstem were mounted sequentially onto slides (96 sections/mouse) and imaged at 10x (Zeiss PlanNeoFluor objective, 10x NA0.3) using a Zeiss AxioImagerM2 fluorescence microscope with an OrcaFlash4.0 camera and Ludl motorized 2axis stage controlled by Neurolucida software (MBF Bioscience, Williston, VT). Each 50 μm section was imaged in 7 μm steps, and the tiled images were then compiled into a single image and collapsed into a single plan using the DeepFocus function in Neurolucida. Analysis of labeled neurons by reconstructing the series of serial sections into a whole brain volume, detecting and marking labeled neurons and registering the location of neurons labeled with tdTomato into the Allen Brain Atlas using NeuroInfo Software^73^. To display the brain wide distribution of cortical neurons providing synaptic inputs to striatal ChAT neurons coordinates of presynaptic rabies labeled cortical neurons registered to the Allen CCF are projected along curved streamlines of the cortex to a flattened mapping of the cerebral cortex^74^. Data were further analyzed in Python and are presented as a percentage of the total labeled cells in each overall region (cortex, thalamus, midbrain, and basal ganglia).

## Acknowledgments

We would like to acknowledge the training and/or support from Jonathan Kuo, Sarah Williams Avram, Vitaly Boyko, Snehashis Roy, and Ted Usdin from the Systems Neuroscience Imaging Resource within the National Institute of Mental Health Intramural Program. We would also like to acknowledge Scott Gerfen (MBF Biosciences) for his assistance with the “flat” cortical maps in Figures 2 and S2. This work was supported by the NIH Center on Compulsive Behaviors (to H.C.G, L.M.A, and V.A.A) and the Intramural Programs of the National Institutes of Mental Health (V.A.A.: ZIAMH002987; C.R.G.: ZIAMH002497), the National Institute on Alcohol Abuse and Alcoholism (V.A.A.: ZIAA000421), and the National Eye Institute (R.J.K.: ZIAEY000511). The contributions of the NIH authors were made as part of their official duties as NIH federal employees, are in compliance with agency policy requirements, and are considered Works of the United States Government. However, the findings and conclusions presented in this paper are those of the authors and do not necessarily reflect the views of the NIH or the U.S. Department of Health and Human Services.

---

**Supplemental Figure 1.**
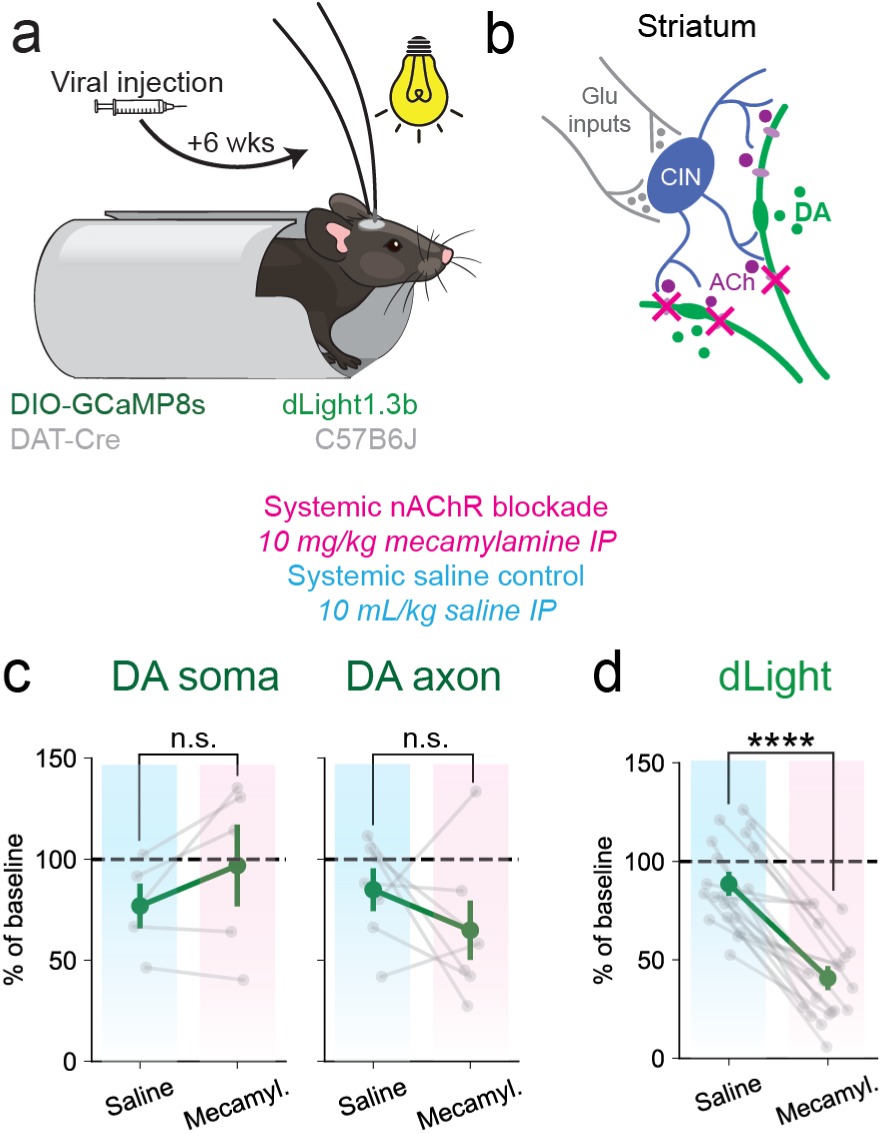
Systemic mecamylamine suppresses striatal cue-evoked dopamine responses. **a.** Experimental setup. DAT-IRES-cre mice expressing DIO-GCaMP8s or wildtype mice expressing dLight1.3b in DMS were head-fixed and presented with white light (500 ms, 5.9 cd/m^2^) and tones (7.5 kHz, 500 ms), spaced roughly every 30 seconds. DAT-GCaMP signals were recorded from both VTA/SNc (somas) and DMS (axons). **b.** Diagram of proposed striatal microcircuitry: cholinergic interneurons (CIN, blue) are driven by glutamatergic inputs (gray), leading to release of acetylcholine (purple), which activates nAChRs on dopamine neuron axons (green) to produce local dopamine release. **c.** Neither intraperitoneal saline injections nor mecamylamine injection affected DA somas or axons. Saline reduced soma responses to 77% of baseline vs. a reduction to 97% by mecamylamine. Saline reduced axonal responses to 85% of baseline vs. a reduction to 65% by mecamylamine (treatment p = 0.26, recording location p = 0.42, interaction p = 0.45; two-way repeated measures ANOVA). **d.** Intraperitoneal mecamylamine injection significantly reduced dLight1.3 responses to the light stimulus. Saline reduced dLight responses to 89 ± 4.8% of baseline (mean ± sem). Mecamylamine reduced dLight responses to 41 ± 4.8% of baseline (treatment p < 0.0001, one-way repeated measures ANOVA).

**Supplemental Figure 2.**
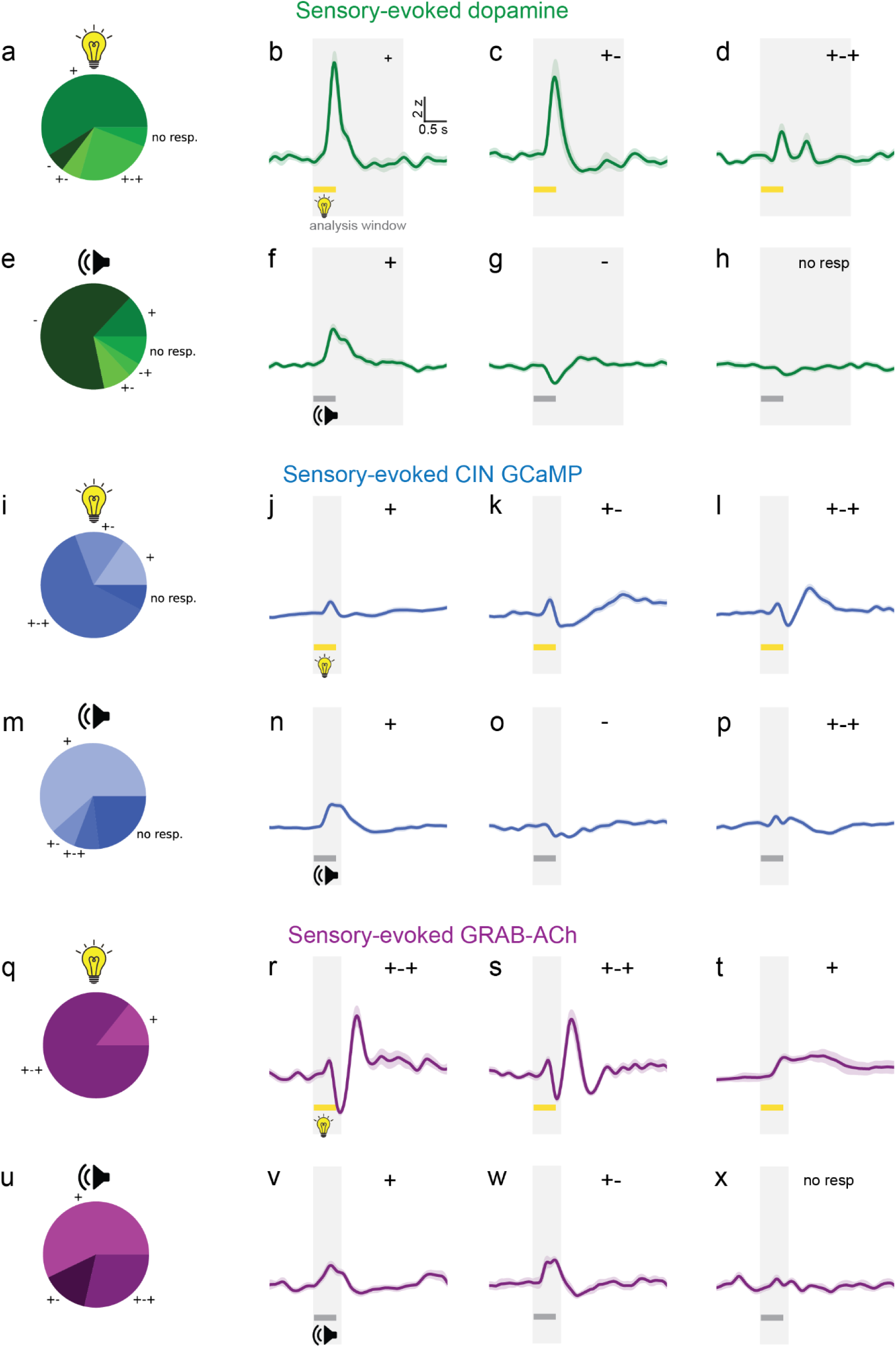
Heterogeneity in cue responses across mice. **a, e.** Percentages of mice that exhibited positive monophasic (+), inverted monophasic (-), biphasic (+-), inverted biphasic (-+), and triphasic dopamine responses to cue and tone flashes (+-+). **b-d, f-h.** Example traces from three different mice, demonstrating heterogeneity in dopamine responses. Gray shaded regions indicate the time window used for initial peak-detection (2 s). **i, m.** Percentages of mice that exhibited positive monophasic (+), biphasic (+-), and triphasic CIN GCaMP responses to cue and tone flashes. **j-k, n-p.** Example traces from three different mice, demonstrating heterogeneity in CIN GCaMP responses. Gray shaded regions indicate the time window used for initial peak-detection (0.6 s). **q, u.** Percentages of mice that exhibited positive monophasic (+), biphasic (+-), and triphasic GRAB-ACh responses to cue and tone flashes. **r-t, v-x.** Example traces from three different mice, demonstrating heterogeneity in acetylcholine responses. Gray shaded regions indicate the time window used for initial peak-detection (0.6 s).

**Supplemental Figure 3.**
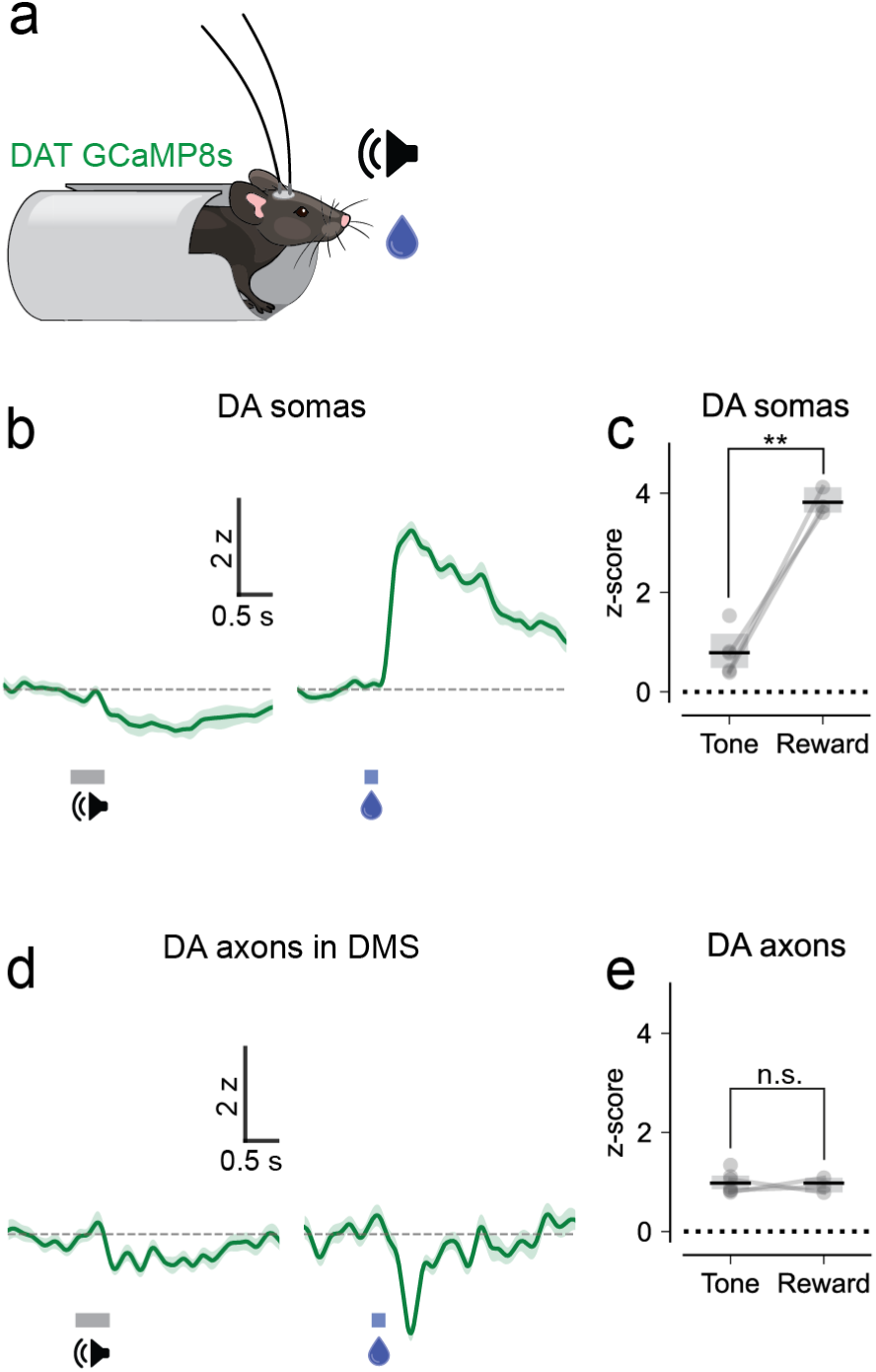
Dopamine neurons respond more strongly to reward than to tone. **a** Water-restricted DAT-IRES-cre mice expressing DIO-GCaMP8s were headfixed and presented with auditory tones and water-drop rewards while fluorescent calcium signals were recorded from midbrain and DMS. **b.** Representative GCaMP8s fluorescence traces from DA somas in response to auditory tones and water reward droplets. **c.** Maximum z-scored GCaMP8s responses across animals (tone, mean 0.79 z [0.48 - 1.1] bootstrap 95% CI; reward, 3.8 z [3.6 - 4.1]; p = 0.005, one-way repeated measures ANOVA). **d.** Representative GCaMP8s fluorescence traces from DA axons in response to auditory tones and water reward droplets. **e.** Maximum z-scored GCaMP8s responses across animals (tone, mean 0.98 z [0.85 - 1.1] bootstrap 95% CI; reward, 0.92 z [0.77 - 1.1]; p = 0.86, one-way repeated measures ANOVA).

**Supplemental Figure 4.**
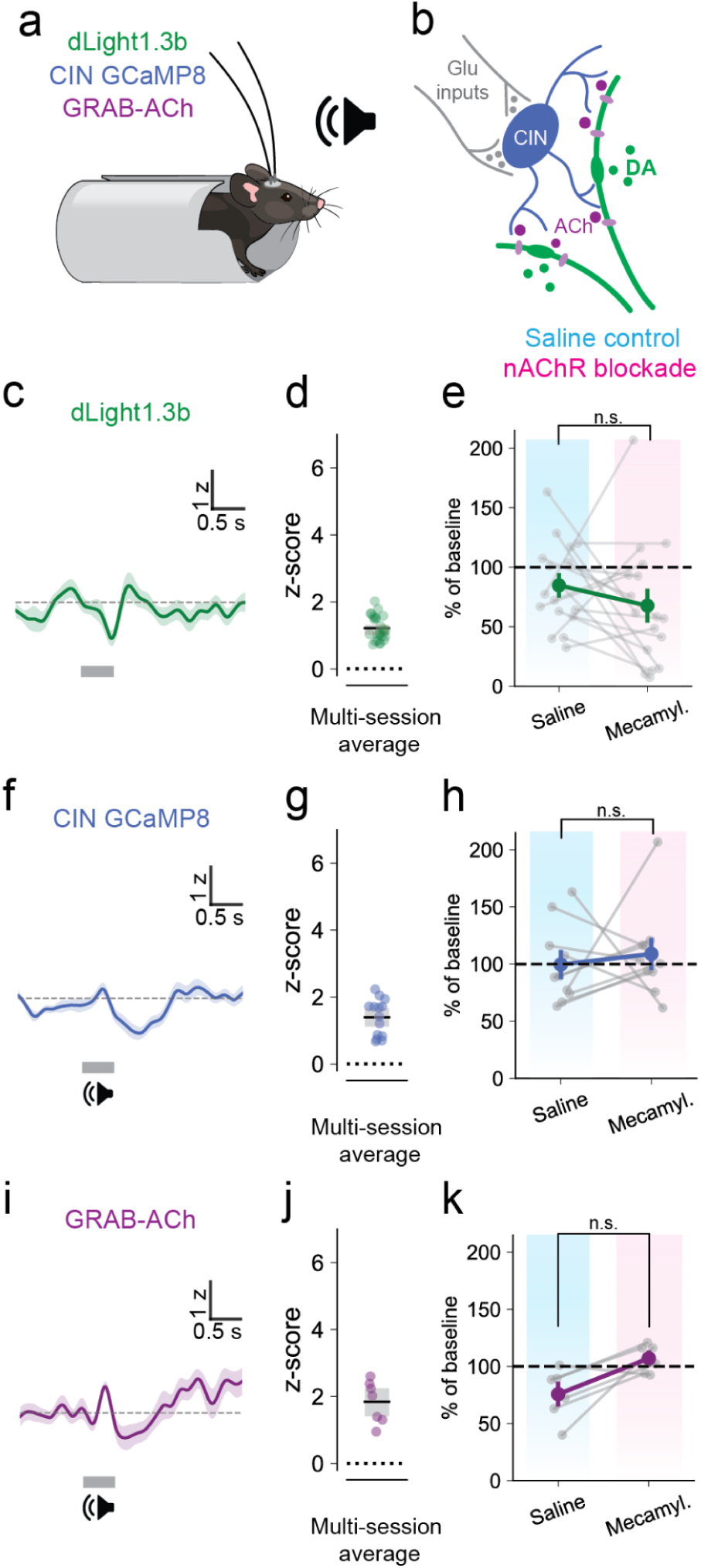
Systemic mecamylamine does not affect tone-evoked CIN or acetylcholine responses. **a-b** Implanted mice expressing dLight1.3b, GCaMP8 in CINs or D1-MSNs, or GRAB-ACh were headfixed and presented with 500 ms tones. Mice were presented with the stimuli before and after receiving an intraperitoneal injection of saline or 10 mg/kg mecamylamine. **c-d.** Representative traces from one animal expressing dLight1.3b in DMS (c) and maximum evoked z-scored responses across animals (d, n = 21 animals, mean response 1.2 z [1.1 - 1.4] bootstrapped 95% CI). **e.** Neither saline nor systemic mecamylamine affected tone-evoked dLight1.3b responses (n = 20 animals; saline mean 85% ± 8.6% SEM; mecamylamine mean 67% ± 12% SEM; p = 0.29, one-way repeated measures ANOVA). **f-g.** Representative traces from one animal expressing GCaMP8s in striatal CINs (f) and maximum evoked z-scored responses across animals (g, n = 12 animals mean response 1.4 z [1.1 - 1.7] bootstrapped 95% CI). **h.** Neither saline nor systemic mecamylamine affected tone-evoked CIN-GCaMP8 responses (n = 11 animals; saline mean 100% ± 11% SEM; mecamylamine mean 89% ± 12% SEM; p = 0.95, one-way repeated measures ANOVA). **i-j.** Representative traces from one animal expressing GRAB-ACh in striatal CINs (f) and maximum evoked z-scored responses across animals (g, mean response 1.8 z [1.4 - 2.3] bootstrapped 95% CI). **k.** Saline reduced tone-evoked GRAB-ACh responses mildly more than mecamylamine (n = 6 animals; saline mean 76% ± 9% SEM; mecamylamine mean 107% ± 5% SEM; p = 0.035, one-way repeated measures ANOVA).

**Supplemental Figure 5.**
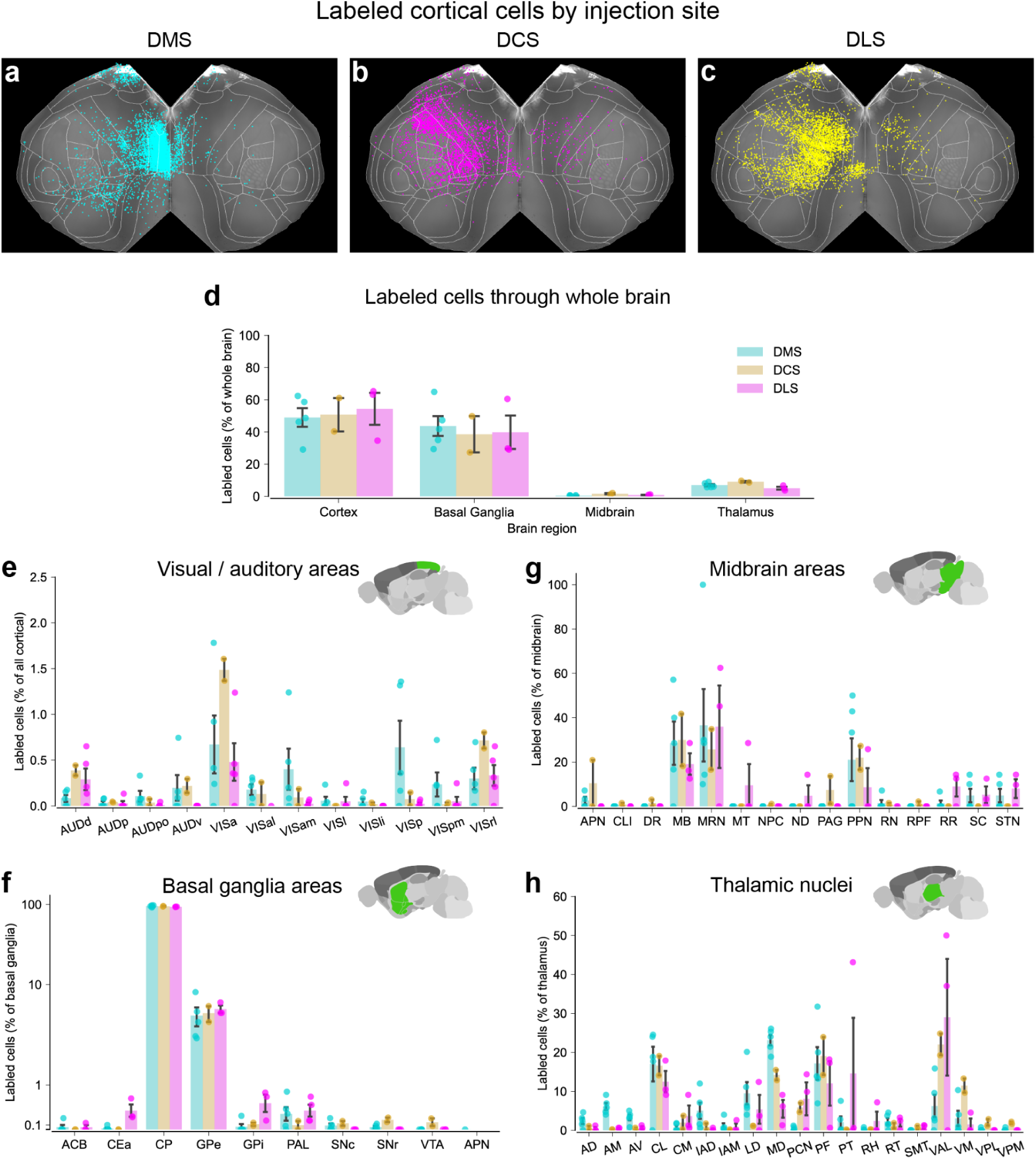
Retrograde rabies tracing of monosynaptic projections to CINs - regional labeling. **a-c** Top-down whole brain views of presynaptic rabies labeled cortical neurons providing inputs to CINs in DMS (cyan, left), DCS (yellow, center), and DLS (magenta, right). **d.** Labeled cells across the cortex, basal ganglia, thalamus, and midbrain as a percentage of all cells labeled throughout the brain. Regardless of injection site, expression was more widespread across the cortex and basal ganglia than in the midbrain or thalamus. **e.** Cells labeled in primary and secondary visual and auditory cortices, as a percentage of all labeled cortical cells. Only a very small percentage of labeled cells (0.5-1.5%) were found in these regions. However, it should be noted that more labeling was found in some secondary visual areas (VISa, VISrl) than in the primary visual cortex. **f.** Percent of labeled cells within basal ganglia subregions, as a percent of all labeled cells in basal ganglia. Note that the y-axis is log-scaled. Most labeled cells were located locally within the striatum (CP, caudate-putamen), with additional labeling of the globus pallidus (GPe, GPi). Lesser expression was found throughout the rest of the basal ganglia (ACB, PAL, SNc, SNr, ACB, VTA, CEa). **g.** Labeled cells within the midbrain, by subregion, as a percentage of all labeled cells in the midbrain. Labeling in the midbrain was sparse; however, most labeling was seen in MRN, PPN, and other unspecified midbrain areas (MB). **h.** Percent of labeled cells within the thalamus by subregion, as a percentage of all labeled cells in the thalamus. Injections across the dorsal striatum produced labeling in CL and PF, while medial injections produced more labeling in MD, and lateral injections produced more labeling in PCN and VAL. For all panels, the area name and abbreviation follow Allen Institute naming conventions (**Table S2**).

**Supplemental Figure 6.**
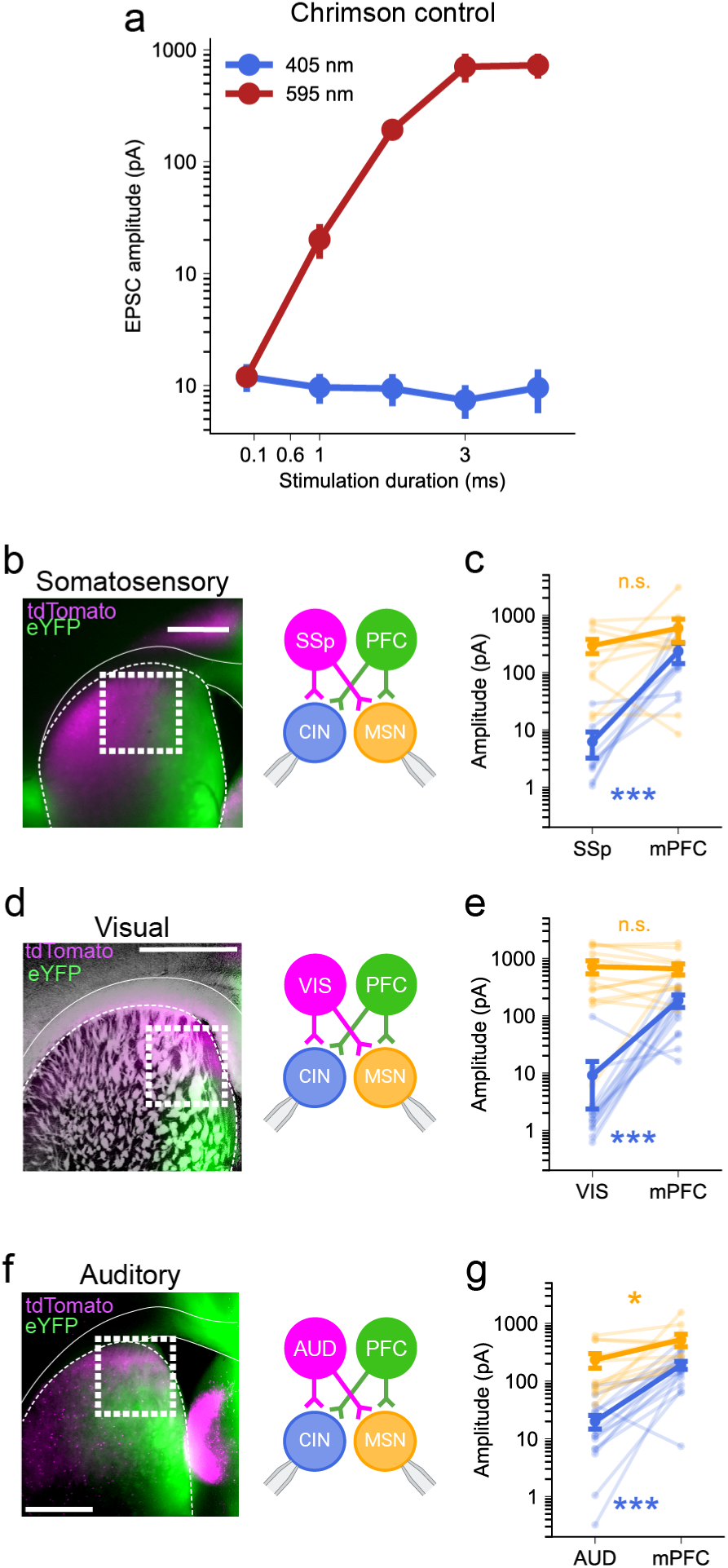
Whole-cell voltage clamp data using dual optogenetic stimulation. To directly compare how single cells respond to input from sensory areas and frontal areas, we expressed in the same animal ChrimsonR-tdT in a sensory area and ChR2-eYFP in the prelimbic cortex. This allowed us to record from striatal cells in areas of overlapping expression and measure synaptic responses evoked by 405 nm violet light to drive ChR2-expressing terminals and 590 nm light to drive ChrimsonR-expressing terminals. The opsin’s selection was made with the intention of minimizing false positive responses when stimulating sensory cortex axons. ChR2 is activated by blue light, which can also spuriously activate ChrimsonR. Red light, however, provides cleaner activation of only ChrimsonR. Therefore, we expressed ChR2 in prelimbic terminals and ChrimsonR in sensory terminals. This way, red light activation of ChrimsonR would exclusively activate sensory terminals. If the expression were swapped, blue light stimulation of sensory terminals would also stimulate the stronger projections from the prelimbic cortex, resulting in false positives. **a.** Effects of 405 nm (off-target) and 595 nm (on-target) stimulation in an animal that only expressed ChrimsonR. 405 nm violet light does not activate ChrimsonR terminals. **b, d, f.** Coronal brain sections from ChAT-tdTomato reporter mice expressing ChR2-eYFP in PL (green) and ChrimsonR-tdTomato in either SSp (a), VISp (c), or AUDp (e, magenta). Dual opsin optogenetic stimulation was used to probe for evoked synaptic responses while recording from striatal CINs (blue) and MSNs (orange) in areas of axonal overlap (white squares). Excitatory synaptic responses were measured using whole-cell voltage-clamp electrophysiology. All white scale bars are 1 mm. **c, e, g.** Mean amplitude of EPSCs recorded from CINs (blue) and MSNs (orange) following optogenetic stimulation of PL and sensory corticostriatal projections. Dark dots represent population mean ± SEM, and light dots represent data from each cell recorded.

**Figure Supplement 7.**
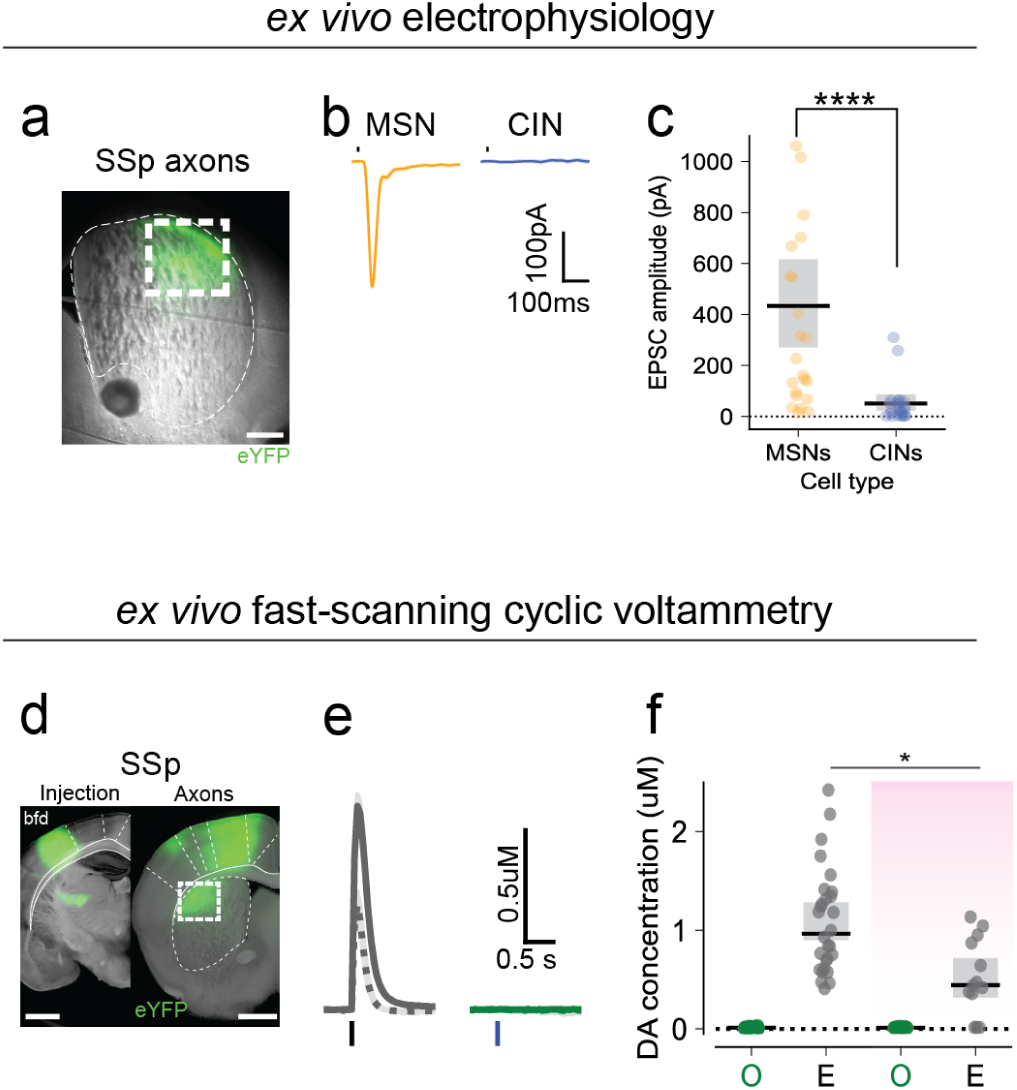
Somatosensory terminals in dorsolateral striatum do not recruit CINs or produce cholinergic-dependent dopamine. Mice with labeled CINs (ChAT-IRES-Cre x tdTomato) were injected with either ChR2-eYFP or ChrimsonR-tdTomato in primary somatosensory cortex (SSp). Brains were extracted and sliced for ex vivo whole-cell patch clamp electrophysiology and fast-scanning cyclic voltammetry following a minimum four-week incubation. **a.** Representative image of ChR2-eYFP labeled SSp projections in striatum. Recordings were conducted within the white box. Scale bar is 1mm. **b.** Representative traces of excitatory postsynaptic currents recorded from an MSN and a CIN in DLS. **c.** Average MSN EPSC amplitudes evoked via optogenetic stimulation of SSp projections (MSN mean 433 pA [270 - 622] bootstrapped 95% CI n = 16 slices from 4 mice; CIN mean 50 pA [22 - 87], n = 14 slices from 5 mice; p < 0.00001, one-way ANOVA). **d.** Left, ChR2-eYFP expression (green) in SSp. Right, ChR2-eYFP expressing projections (green) from SSp in dorsomedial striatum. The white box shows the site of dopamine recordings. Scale bars are 1 mm. **e.** Single-slice average of dopamine transients measured with fast-scanning cyclic voltammetry (FSCV) and evoked by either optogenetic stimulation of corticostriatal projections (green) or electrical stimulation (gray) in the same recording site. Dashed lines show the remaining dopamine responses after washing in a nicotinic receptor blocker (1 μM DHβE). Traces are mean and shaded areas are ± SEM). **f.** Average amplitudes of dopamine transient in DLS elicited by electrical stimulation (gray bars, in ACSF mean 1.09 uM [0.90 - 1.3] bootstrapped 95% CI; in DHBE mean 0.52 uM [0.32 - 0.72]; ACSF vs DHBE p = 0.005, Mann-whitney U) and optogenetic stimulation (green bars, in ACSF mean 0.013 uM [0.011 - 0.016]; in DHBE mean 0.014 [0.011 - 0.017]) before and after DHβE bath application (pink shade). Dots represent data from individual slices. Lines and shaded bars are mean ± sem.

**Supplemental Figure 8:**
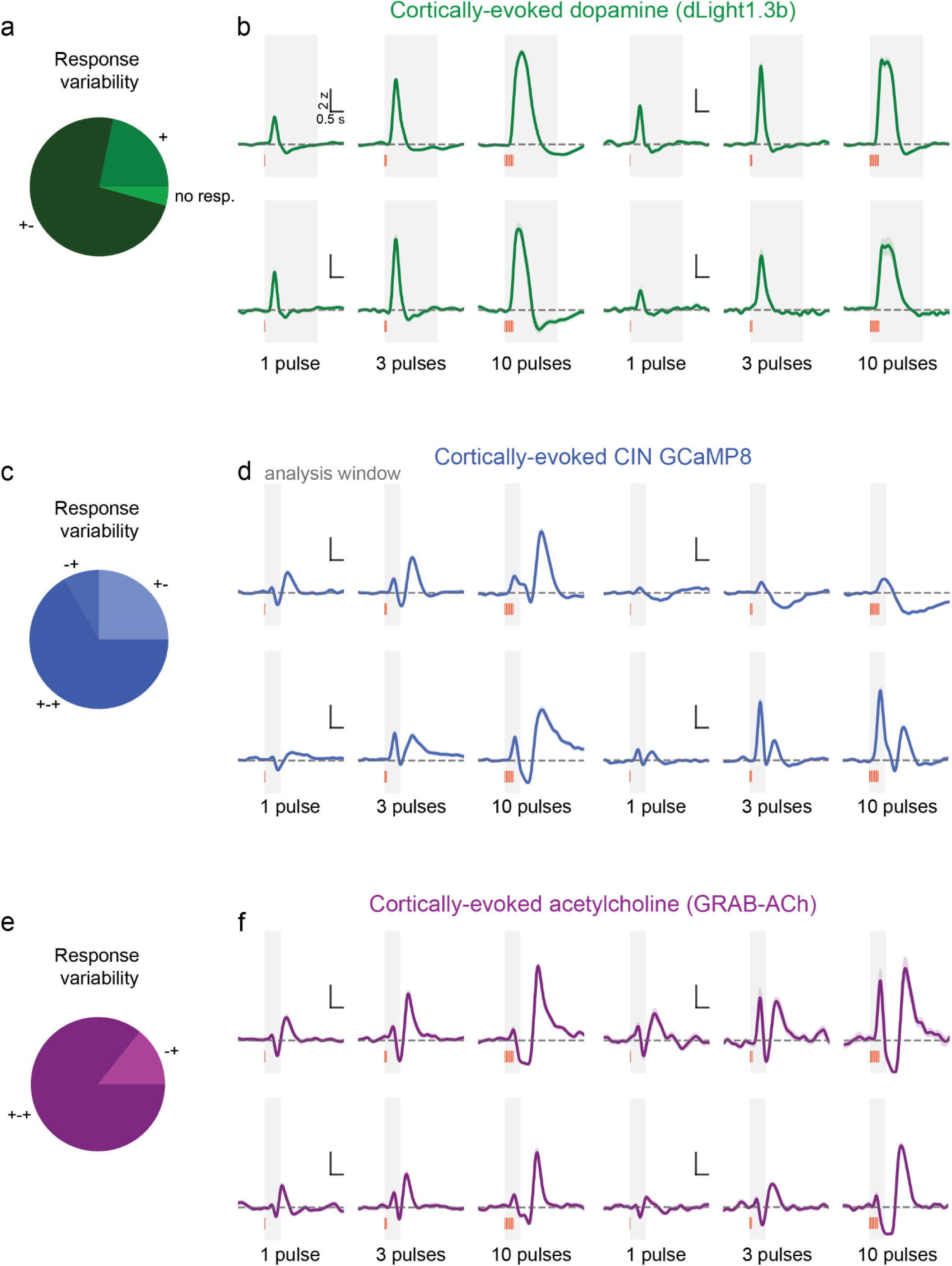
Heterogeneity in cortically-evoked responses. **a** Percentages of animals exhibiting monophasic (+), biphasic (+-) or no dopamine response to cortical stimulation. **b.** Example traces of cortically-evoked dopamine responses from four animals, demonstrating heterogeneity in the second phase of the response. Gray shaded region represents the 2 s time window used for analysis. **c,e.** Percentages of animals exhibiting biphasic (+-), inverse biphasic (-+), or triphasic CIN GCaMP © or GRAB-ACh (e) responses to cortical stimulation. **d, f.** Example traces of cortically-evoked CIN GCaMP (d) or GRAB-ACh (f) responses from four animals, demonstrating heterogeneity across all three phases of the response. Because of this heterogeneity, analysis was limited to 0.6s after stimulation, which consistently captured the first phase of the response.

**Supplemental Figure 9:**
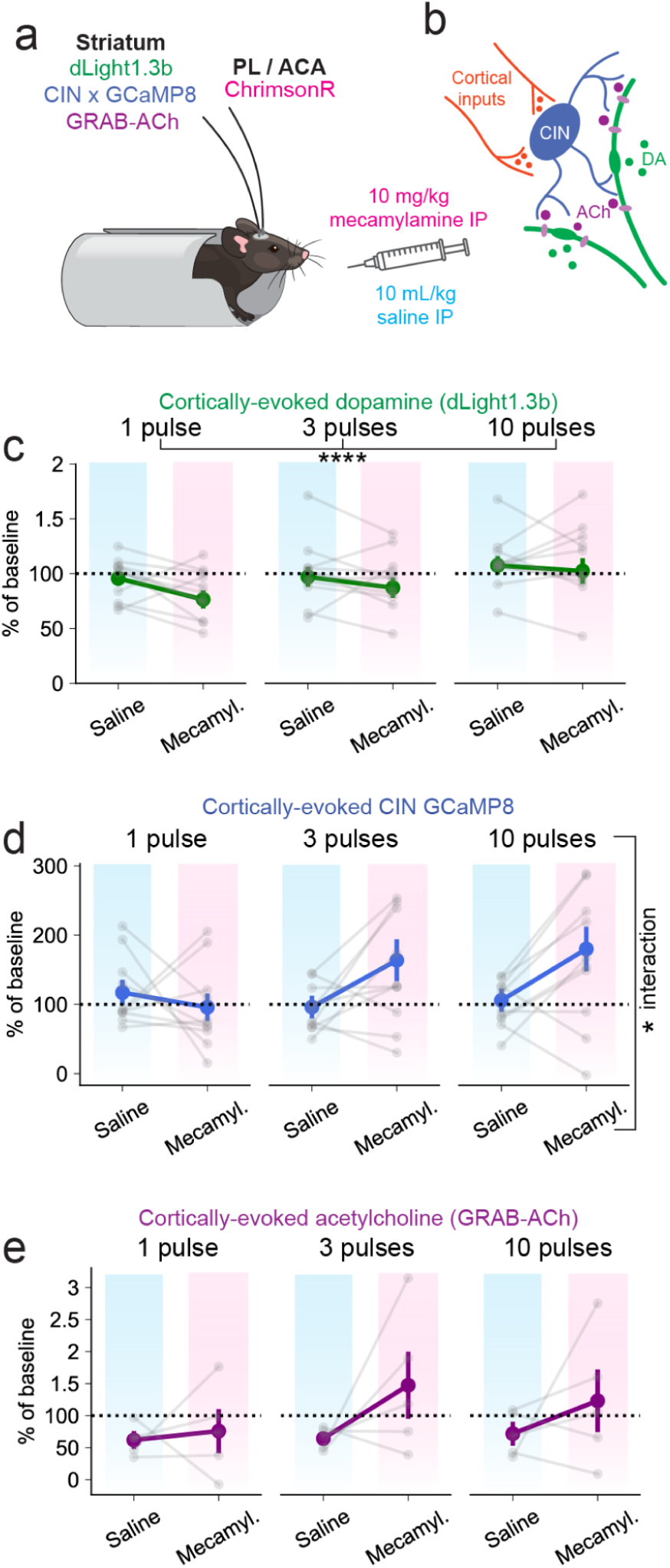
Systemic saline and mecamylamine do not significantly reduce optogenetically evoked responses. **a-b** Implanted mice expressing dLight1.3b, GCaMP8 in CINs, or GRAB-ACh in DMS were headfixed and received optogenetic stimulation of PL / ACA striatal terminals (see Figure 6, Methods). Recordings were conducted before and after injecting the animal intraperitoneally with 10 mL / kg sterile saline (vehicle) or 10 mg / kg mecamylamine. **c.** Systemic saline and mecamylamine had limited effects on cortically-evoked dopamine responses, though there was an effect of stimulation pattern (n = 11 mice, treatment p = 0.31, stimulation p = 0.00007, interaction p = 0.18, two-way repeated measures ANOVA). **d.** Systemic saline and mecamylamine had no effect on cortically-evoked CIN GCaMP responses, though there was an effect of the interaction between treatment and stimulation pattern (n = 10 animals, treatment p = 0.13, stimulation p = 0.48, interaction p = 0.048, two-way repeated measures ANOVA). **e.** Systemic saline and mecamylamine had no effect on cortically-evoked GRAB-ACh responses (n = 5 animals, treatment p = 0.08, stimulation p = 0.57, interaction p = 0.55, two-way repeated measures ANOVA). In all plots, individual gray lines represent data from one animal. Dark lines represent the mean and sem across all animals.

**Supplemental Figure 10:**
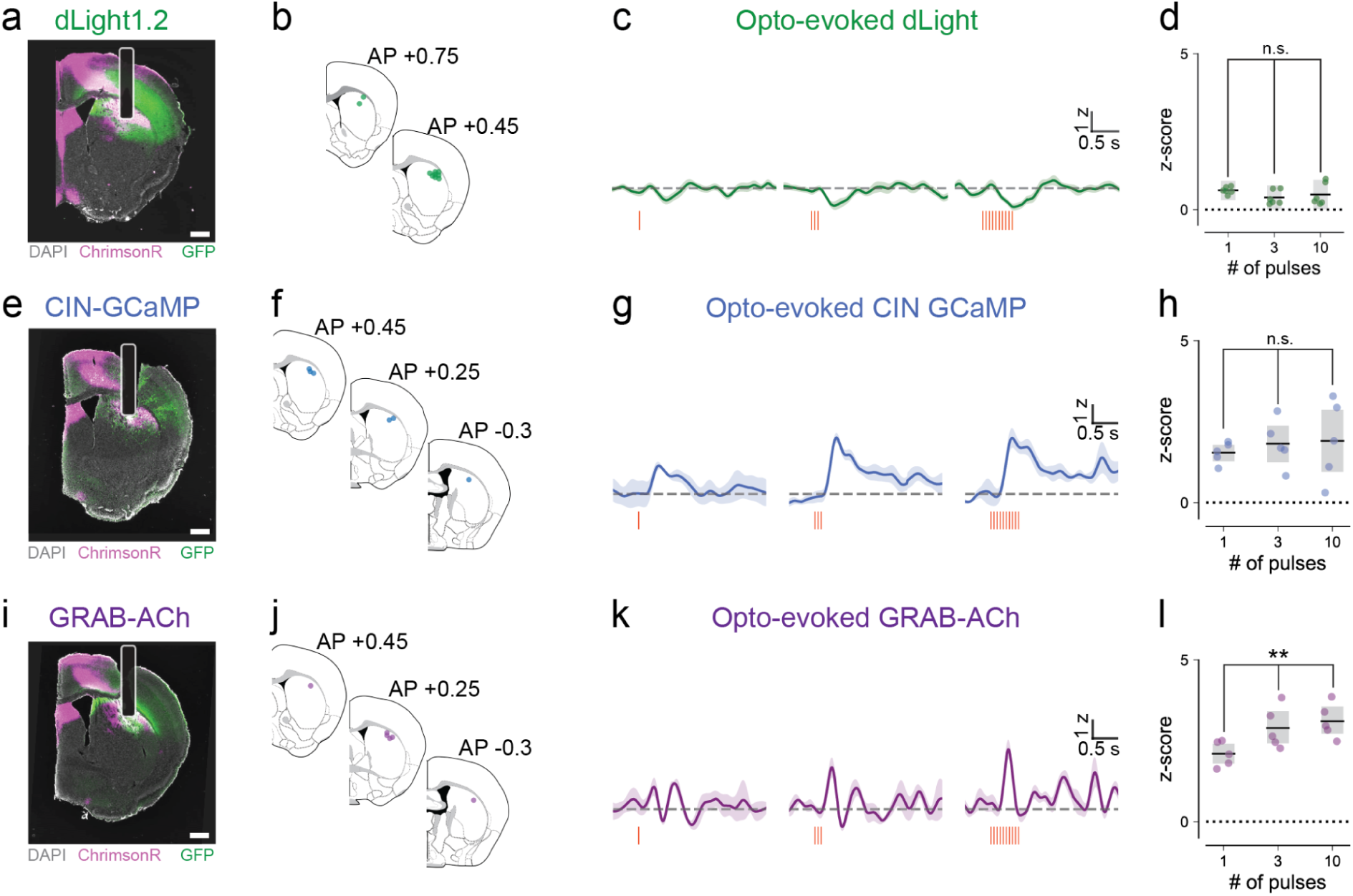
Somatosensory cortex drives CINs and acetylcholine release in dorsolateral striatum, but not dopamine. **a,e,i** Images from ChAT-IRES-cre or wildtype mice were injected with ChrimsonR in primary somatosensory cortex and with dLight1.2, DIO-GCaMP8s, or GRAB-ACh in dorsolateral striatum (green). Animals were implanted with a headpost and bilateral fiber optic cannula in DLS, allowing for stimulation of SSp terminals (magenta). Scale bars are 0.5mm. **b, f, j.** Fiber placements in DLS, n = 6 animals per sensor. Animals expressing dLight1.3b were implanted bilaterally. The average of the two hemispheres is presented as one datapoint. **c, g, k.** Representative traces from three animals expressing dLight1.3b, GCaMP8s in CINs, or GRAB-ACh in DLS. Optogenetic stimulation protocol included a single 10ms pulse of 590 nm red light, trains of 3 pulses at 20 Hz, and trains of 10 pulses at 20 Hz, with each stimulation occuring 30s apart. **d, h, i.** Maximum z-scored optogenetically-evoked responses across animals expressing dLight1.3b (d), CIN GCaMP (h), and GRAB-ACh (i). Stimulation intensity did not affect dLight1.3b or CIN GCaMP (dLight p = 0.07, CIN GCaMP p = 0.48, one-way repeated measures ANOVA), though it did have an effect on GRAB-ACh responses (p = 0.0004, one-way repeated measures ANOVA).

**Supplemental Table 1:** Fiber photometry animals & viral expression.

| Mouse | Sex | Age (m) | Genotype | Left Hem. | Right Hem. | Chrimson |
| --- | --- | --- | --- | --- | --- | --- |
| Ch-18 | F | 9 | WT | dLight1.2b | dLight1.2b | SSp |
| Ch-25 | M | 9 | WT | dLight1.2b | dLight1.2b | SSp |
| Ch-26 | M | 9 | WT | dLight1.2b | dLight1.2b | SSp |
| HG-01 | M | 6 | WT | dLight1.2b | dLight1.2b | SSp |
| HG-02 | M | 6 | WT | dLight1.2b | dLight1.2b | SSp |
| HG-03 | M | 6 | WT | dLight1.2b | dLight1.2b | SSp |
| HG-04 | M | 4 | ChAT-Cre |  | GCaMP8s | PL |
| HG-05 | M | 6 | ChAT-Cre |  | GCaMP8s | PL |
| HG-06 | M | 7 | ChAT-Cre | ACh3.0 | GCaMP8s | PL |
| HG-08 | F | 3 | ChAT-Cre | ACh3.0 | GCaMP8s | PL |
| HG-09 | F | 3 | ChAT-Cre | ACh3.0 | GCaMP8s | PL |
| HG-10 | F | 3 | ChAT-Cre | ACh3.0 | GCaMP8s | PL |
| HG-11 | F | 6 | ChAT-Cre | GCaMP8f | dLight1.3b | PL |
| HG-12 | F | 6 | ChAT-Cre | GCaMP8f | dLight1.3b | PL |
| HG-14 | M | 6 | ChAT-Cre | GCaMP8f | dLight1.3b | PL |
| HG-15 | F | 3 | ChAT-Cre | GCaMP8f | GCaMP8f | ACA |
| HG-16 | F | 3 | ChAT-Cre | GCaMP8f | GCaMP8f | ACA |
| HG-17 | M | 3 | ChAT-Cre | GCaMP8f | dLight1.3b | ACA |
| HG-18 | M | 5 | WT | ACh3.0 | dLight1.3b | ACA |
| HG-19 | F | 5 | WT | ACh3.0 | dLight1.3b | ACA |
| HG-20 | M | 4 | WT | ACh3.0 | dLight1.3b | ACA |
| HG-21 | F | 5 | DAT-Cre | GCaMP8s |  | PL |
| HG-22 | M | 6 | DAT-Cre | GCaMP8s |  | PL |
| HG-23 | M | 6 | WT | dLight1.3b |  | PL |
| HG-24 | M | 6 | WT | dLight1.3b* |  | PL |
| HG-25 | M | 7 | ChAT-Cre | GCaMP8s | ACh3.0 | SSp |
| HG-26 | M | 7 | ChAT-Cre | GCaMP8s | ACh3.0 | SSp |
| HG-27 | M | 7 | ChAT-Cre | GCaMP8s | ACh3.0 | SSp |
| HG-28 | M | 3 | DAT-cre | GCaMP8s |  | PL |
| HG-29 | M | 3 | DAT-cre | GCaMP8s |  | PL |
| HG-30 | M | 3 | DAT-cre | GCaMP8s |  | PL |
| HG-31 | M | 3 | ChAT-Cre | GCaMP8s | ACh3.0 | SSp |
| HG-32 | M | 3 | ChAT-Cre | GCaMP8s | ACh3.0 | SSp |
| HG-33 | M | 3 | ChAT-Cre | GCaMP8s | ACh3.0 | SSp |

**Supplemental Table 1 continued: Fiber photometry animals & viral expression**
| Mouse | Sex | Age (m) | Genotype | Left Hem. | Right Hem. | Chrimson |
| --- | --- | --- | --- | --- | --- | --- |
| HG-34 | F | 3 | DAT-cre | GCaMP8s |  | PL |
| HG-35 | F | 3 | DAT-cre | GCaMP8s |  | PL |
| HG-39 | M | 3 | WT | dLight1.3b* |  | PL |
| HG-40 | M | 3 | WT | dLight1.3b* |  | PL |
| HG-41 | M | 3 | WT | dLight1.3b* |  | PL |
| HG-42 | F | 3 | WT | dLight1.3b* |  | PL |
| HG-43 | F | 3 | WT | dLight1.3b* |  | PL |
| HG-44 | F | 3 | WT | dLight1.3b* |  | PL |
| RR-03 | M | 4 | WT | dLight1.3b | dLight1.3b | PL |
| RR-05 | M | 4 | WT | dLight1.3b | dLight1.3b | PL |
| RR-07 | M | 4 | WT | dLight1.3b | dLight1.3b | PL |
| RR-09 | M | 4 | WT | dLight1.3b | dLight1.3b | PL |
| LA-11 | M | 7 | D1-Cre | GCaMP8s | GCaMP8s |  |
| LA-12 | M | 7 | D1-Cre | GCaMP8s | GCaMP8s |  |
| LA-13 | M | 7 | D1-Cre | GCaMP8s | GCaMP8s |  |
| LA-15 | M | 9 | D1-Cre | GCaMP8s | GCaMP8s |  |
| LA-16 | M | 9 | D1-Cre | GCaMP8s | GCaMP8s |  |
| LA-18 | F | 8 | D1-Cre | dLight1.3b | GCaMP8s |  |
| LA-19 | F | 8 | D1-Cre | dLight1.3b | GCaMP8s |  |
| LA-20 | F | 8 | D1-Cre | GCaMP8s | dLight1.3b |  |
| LA-28 | M | 6 | D1-Cre | PFC axon ipsi | PFC axon contra |  |
| LA-29 | M | 6 | D1-Cre | PFC axon ipsi | PFC axon contra |  |
| LA-30 | F | 6 | D1-Cre | PFC axon ipsi | PFC axon contra |  |
| LA-33 | M | 6 | Pdyn-Cre | PFC axon ipsi | PFC axon contra |  |
| LA-34 | F | 6 | Pdyn-Cre | PFC axon ipsi | PFC axon contra |  |
| LA-35 | F | 6 | Pdyn-Cre | PFC axon ipsi | PFC axon contra |  |
| LA-36 | F | 6 | Pdyn-Cre | PFC axon ipsi | PFC axon contra |  |

\* optofluid cannula

**Supplemental Table 2:**
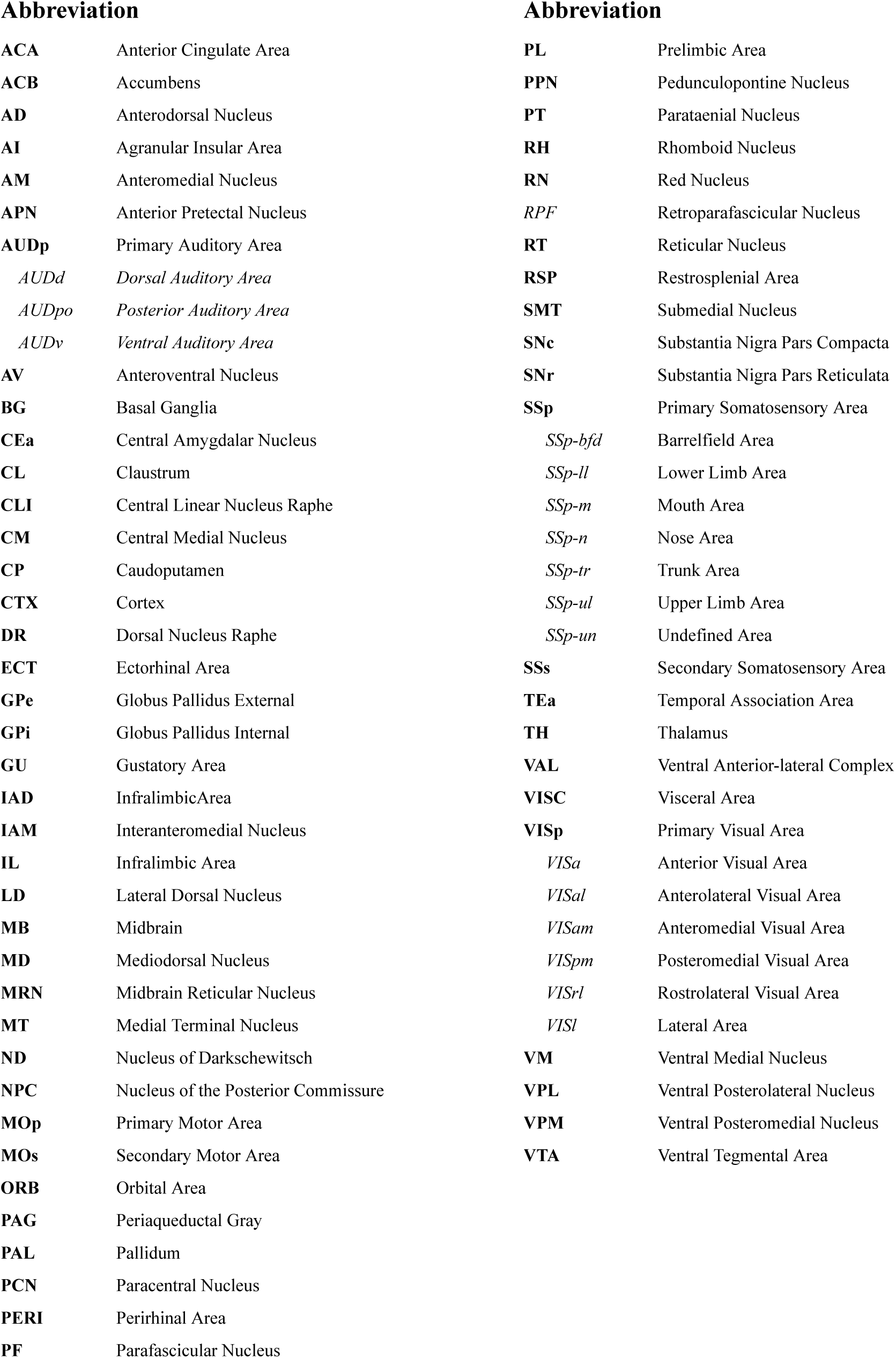
Allen Institute Brain Region Nomenclature.

## Notes

### Competing Interest Statement

The authors have declared no competing interest.

### Summary of Updates

New in vivo data, including local infusions, midbrain GCaMP recordings, DLS recordings, and auditory responses

